# Nuclear NAD^+^-biosynthetic enzyme NMNAT1 facilitates survival of developing retinal neurons

**DOI:** 10.1101/2021.05.05.442836

**Authors:** David Sokolov, Emily Sechrest, Yekai Wang, Connor Nevin, Jianhai Du, Saravanan Kolandaivelu

## Abstract

Despite mounting evidence that the mammalian retina is exceptionally reliant on proper NAD^+^ homeostasis for health and function, the specific roles of subcellular NAD^+^ pools in retinal development, maintenance, and disease remain obscure. Here, we show that deletion of the nuclear-localized NAD^+^ synthase nicotinamide mononucleotide adenylyltransferase-1 (NMNAT1) in the developing murine retina causes early and severe degeneration of photoreceptors and select inner retinal neurons via multiple distinct cell death pathways. This severe phenotype is associated with disruptions to retinal central carbon metabolism, purine nucleotide synthesis, and amino acid pathways. Furthermore, large-scale transcriptomics reveals dysregulation of a collection of photoreceptor and synapse-specific genes in NMNAT1 knockout retinas prior to detectable morphological or metabolic alterations. Collectively, our study reveals previously unrecognized complexity in NMNAT1-associated retinal degeneration and suggests a yet-undescribed role for NMNAT1 in gene regulation during photoreceptor terminal differentiation.

## INTRODUCTION

Nicotinamide adenine dinucleotide (NAD^+^) is a ubiquitous cellular metabolite with an ever-expanding palette of biological functions across all kingdoms of life. In addition to serving a central role in redox metabolism as an electron shuttle, NAD^+^ has well-defined roles as a substrate for a host of enzymes including sirtuins (SIRTs), mono- and poly- ADP-ribose polymerases (PARPs), and NAD^+^ glycohydrolases (CD38, CD157, and SARM1). Collectively, these roles implicate NAD^+^ metabolism in phenomena as diverse as aging, cell proliferation, immunity, neurodegeneration, differentiation, and development (Houtkooper et al., 2010; Canto et al., 2015; Nikiforov et al., 2015; Cambronne and Kraus, 2020; Navas and Carnero, 2021). A relatively recent advance in the field is the notion of compartmentalized NAD^+^ metabolism—that regulation of NAD^+^ in distinct subcellular compartments dictates function in diverse manners (Canto et al., 2015; Nikiforov et al., 2015; Cambronne and Kraus, 2020; Navas and Carnero, 2021). While many aspects of this compartmentalization remain to be explored, it is now known that spatiotemporal NAD^+^ regulation plays prominent roles in processes including axon degeneration, circadian regulation, and adipogenesis (Cambronne and Kraus, 2020).

Among mammalian tissues, the retina appears particularly reliant on proper NAD^+^ homeostasis for survival and function. This is suggested by associations between retinal NAD^+^ deficiency and pathology in diverse models of retinal dysfunction (Lin et al., 2016; Williams et al., 2017) as well as multiple mutations to NAD^+^- or NADP^+^-utilizing enzymes which cause blindness in humans (Bowne et al., 2006; Aleman et al., 2018; Bennett et al., 2020). Among these enzymes is nicotinamide adenylyltransferase 1 (NMNAT1), a highly conserved, nuclear-localized protein which catalyzes the adenylation of nicotinamide mononucleotide (NMN) or nicotinic acid mononucleotide (NaMN) to form NAD^+^, the convergent step of all mammalian NAD^+^ biosynthetic pathways (Nikiforov et al., 2015). To date, over 30 NMNAT1 mutations have been linked to the severe blinding diseases Leber congenital amaurosis type 9 (LCA9) and related cone-rod dystrophy (Perrault et al., 2012; Falk et al., 2012; Chiang et al., 2012; Koenekoop et al., 2012; Coppieters et al., 2015; Nash et al., 2018). Although NMNAT1 is ubiquitously expressed, and many of these mutations reduce NMNAT1 catalytic activity or stress-associated stability (Falk et al., 2012; Koenekoop et al., 2012; Sasaki et al., 2015), patients with these disorders rarely report extra-ocular phenotypes, a puzzling observation which is recapitulated by two LCA-NMNAT1 mutant mouse models (Greenwald et al., 2016). Further puzzling is the existence of two other NMNAT isoforms (Golgi-associated NMNAT2 and mitochondrial NMNAT3), which are detectable in the retina (Kuribayashi et al., 2018) but have not been linked to blindness. Importantly, while a crucial role for retinal NAD^+^ was recently described through characterization of mice conditionally lacking the NAD^+^ pathway enzyme NAMPT in photoreceptors (Lin et al., 2016), the significance of nuclear-synthesized NAD^+^ in vision— suggested by the fact that NMNAT1 is the only NAD^+^-pathway enzyme to date linked to blindness—remains poorly understood.

Current results point to multiple, potentially distinct roles for NMNAT1 in the retina—*ex vivo* studies suggest that NMNAT1 supports sirtuin function to facilitate the survival of retinal progenitor cells (Kuribayashi et al., 2018), while ablation of NMNAT1 in mature mice results in rapid death of photoreceptors mediated by the neurodegenerative NADase SARM1 (Sasaki et al., 2020a). Global deletion of NMNAT1 in mice is embryonically lethal (Conforti et al., 2011), suggesting non-redundant roles for nuclear NAD^+^ synthesis during development. Consistent with this notion, pan-retinal NMNAT1 deletion is shown to cause rapid and severe retinal degeneration in mice shortly after birth (Wang et al., 2017; Eblimit et al., 2018). While these studies suggest diverse functions of retinal NMNAT1 beyond its canonical role in redox metabolism, the degree to which these functions overlap—as well as the mechanistic basis for the severity of NMNAT1-associated retinal dystrophy in animal models and patients—have not been comprehensively explored.

In this study, we investigate the roles of NMNAT1-mediated NAD^+^ metabolism in the retina by generating and characterizing a retina-specific NMNAT1 knockout mouse model. Utilizing histological and transcriptomic approaches, we demonstrate that NMNAT1 deletion causes severe and progressive retinal degeneration affecting specific retinal cell types beyond photoreceptors, and that this severe degeneration likely results from activation of multiple distinct cell death pathways. Comprehensive metabolomics analysis reveals specific metabolic defects in NMNAT1 knockout retinas and suggests impaired central carbon, purine nucleotide, and amino acid metabolism as a cause for severe degeneration. Furthermore, concomitant transcriptomics analyses of knockout retinas reveal a cluster of photoreceptor and synapse-specific genes which are downregulated before the onset of degeneration, suggesting a yet-undescribed role for NMNAT1 in gene regulation during late-stage retinal development. Overall, our results reveal a previously unappreciated complexity in NMNAT1-associated retinal degeneration, provide possible explanations for the retina-specific manifestations of NMNAT1 deficiency, and lay the foundation for further study of compartmentalized NAD^+^ metabolism in vision.

## RESULTS

### Generation and validation of NMNAT1 conditional knockout mouse model

To establish a retinal-specific NMNAT1 knockout model, we crossed mice homozygous for a loxP-targeted *Nmnat1* locus (*Nmnat1*^fl/fl^) with transgenic mice expressing Cre recombinase under a *Six3* promoter (*Nmnat1*^wt/wt^ *Six3-Cre*), which activates throughout the retina around embryonic day 9.5 (E9.5) and shows robust activity by E12.5 (Furuta et al., 2000). After several crosses, mice inheriting *Six3-Cre* and a floxed *Nmnat1* locus (*Nmnat1*^fl/fl^ *Six3-Cre*, hereafter referred to as “knockouts”) exhibit Cre-mediated excision of the first two exons of *Nmnat1*—which contain important substrate binding domains—in the embryonic retina (Figure 1A). We determined that retinal *Nmnat1* expression in postnatal day 4 (P4) knockout mice was reduced by 75.6% compared to littermate controls (Figure 1B). We further verified that retinal NMNAT1 protein levels were drastically reduced in P0 knockout mice using a custom-made polyclonal antibody against NMNAT1 (Figure 1C and **Figure 1—Supplement 1**). Finally, we confirmed that embryonic *Six3-Cre* expression alone does not cause gross retinal abnormalities by staining for several well-characterized cell type markers in mature *Nmnat1*^wt/wt^ *Six3-Cre* retinas and littermate controls (**Figure 2—Supplement 1**; markers discussed below).

**Figure 1.**
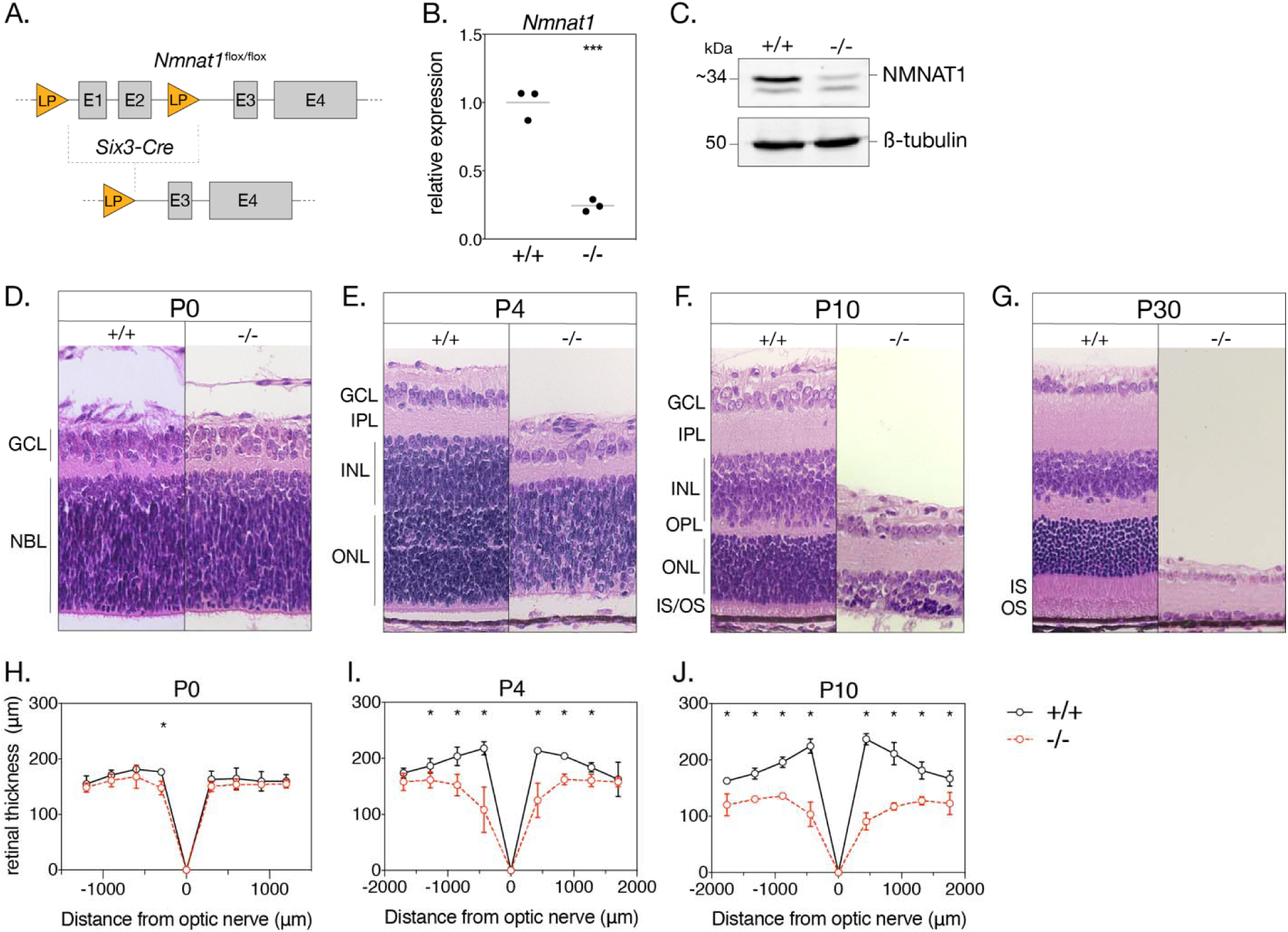
Loss of NMNAT1 leads to early and severe retinal degeneration. **(A**) Schematic depicting retina specific *Six3-Cre* mediated excision of a segment of the *Nmnat1* gene. (**B)** Relative *Nmnat1* expression in retina from P4 knockout (-/-) and littermate control (+/+) mice as assessed by RT-qPCR (grey bars represent mean, *\*\*\**p<0.0005 using Student’s t-test, n=3 biological replicates). (**C)** Representative western blot showing levels of NMNAT1 and β-tubulin loading control in retinal lysate from P0 knockout and control mice. (**D-G**) Representative H&E-stained retinal cross sections from knockout and control mice at indicated ages. (**H-J**) Spider plots depicting mean retinal thickness at P0, P4, and P10. Data are represented as mean ± SD. *p<0.05 using Student’s t-test. Abbreviations: LP, loxP site; E1-4, exon 1-4; P, postnatal day; GCL, ganglion cell layer; NBL, neuroblastic layer; IPL, inner plexiform layer; OPL, outer plexiform layer; INL, inner nuclear layer; ONL, outer nuclear layer; IS/OS, photoreceptor inner segment/outer segment layer.

### Early-onset and severe morphological defects in the NMNAT1-null retina

As a first step towards characterizing the effects of NMNAT1 ablation on the retina, we performed retinal histology using hematoxylin and eosin (H&E) staining. H&E-stained retinal cross sections from P0 knockout and control mice reveal no obvious morphological differences (Figure 1D, H); however, by P4, knockout retina are markedly thinner than controls and exhibit disrupted lamination and evidence of large-scale cell death in the inner and outer nuclear layers (Figure 1E). Degeneration is most severe in the central retina, with a ∼45% reduction in central retinal thickness (but unaffected peripheral retinal thickness) in P4 knockout mice (Figure 1I). By P10, knockouts show a ∼62% reduction in central retinal thickness and ∼27% reduction in peripheral retinal thickness compared to controls (Figure 1J). Degeneration of the entire inner and outer nuclear layers is nearly complete by P30 (Figure 1G), while remaining inner retinal structures persist until approximately P60 (data not shown). Proper segregation of inner and outer retinal neurons appears disrupted in P4 knockouts, but this segregation is established in P10 knockouts despite severe degeneration (Figure 1F). Interestingly, formation of the outer plexiform layer (containing photoreceptor and bipolar neuron synaptic structures) appears disrupted in P4 and P10 knockout retinas (Figure 1E, F).

### NMNAT1 loss affects survival of retinal bipolar and amacrine neurons

Histological examination suggests severe photoreceptor degeneration in NMNAT1 knockout retinas but also indicates loss of specific inner retinal neuron populations. To further characterize these effects, we quantified populations of several major inner retinal cell types in our knockout by staining retinal sections with well-characterized antibody markers: retinal ganglion cells were identified by labelling for brain-specific homeobox/POU domain protein 3A (BRN3A), amacrine cells by labelling for calretinin (CALR), and bipolar cells by labelling for Ceh-10 homeodomain-containing homolog (CHX10) (**Supplemental Table 2**). We performed this analysis at P4 and P10—representing early and late stages of degeneration, respectively—revealing an interesting cell type-dependent sensitivity to NMNAT1 loss (Figure 2). At both tested ages, relative numbers of retinal ganglion cells are not significantly different between knockout and control retina (Figure 2B-E, P), while numbers of amacrine cells are unchanged at P4 but reduced by ∼51% in P10 knockout retinas (Figure 2G-J, Q). Numbers of bipolar cells are similarly unchanged at P4 but reduced by ∼75% in P10 knockout retinas (Figure 2L-O, R). These results identify retinal bipolar and amacrine neurons as targets of NMNAT1-associated degeneration and suggest that bipolar neurons are more sensitive to NMNAT1 loss than amacrine neurons. Interestingly, while retinal ganglion cells do eventually degenerate at timepoints past P30 (data not shown), they appear largely agnostic to NMNAT1 loss in the young postnatal retina.

**Figure 2.**
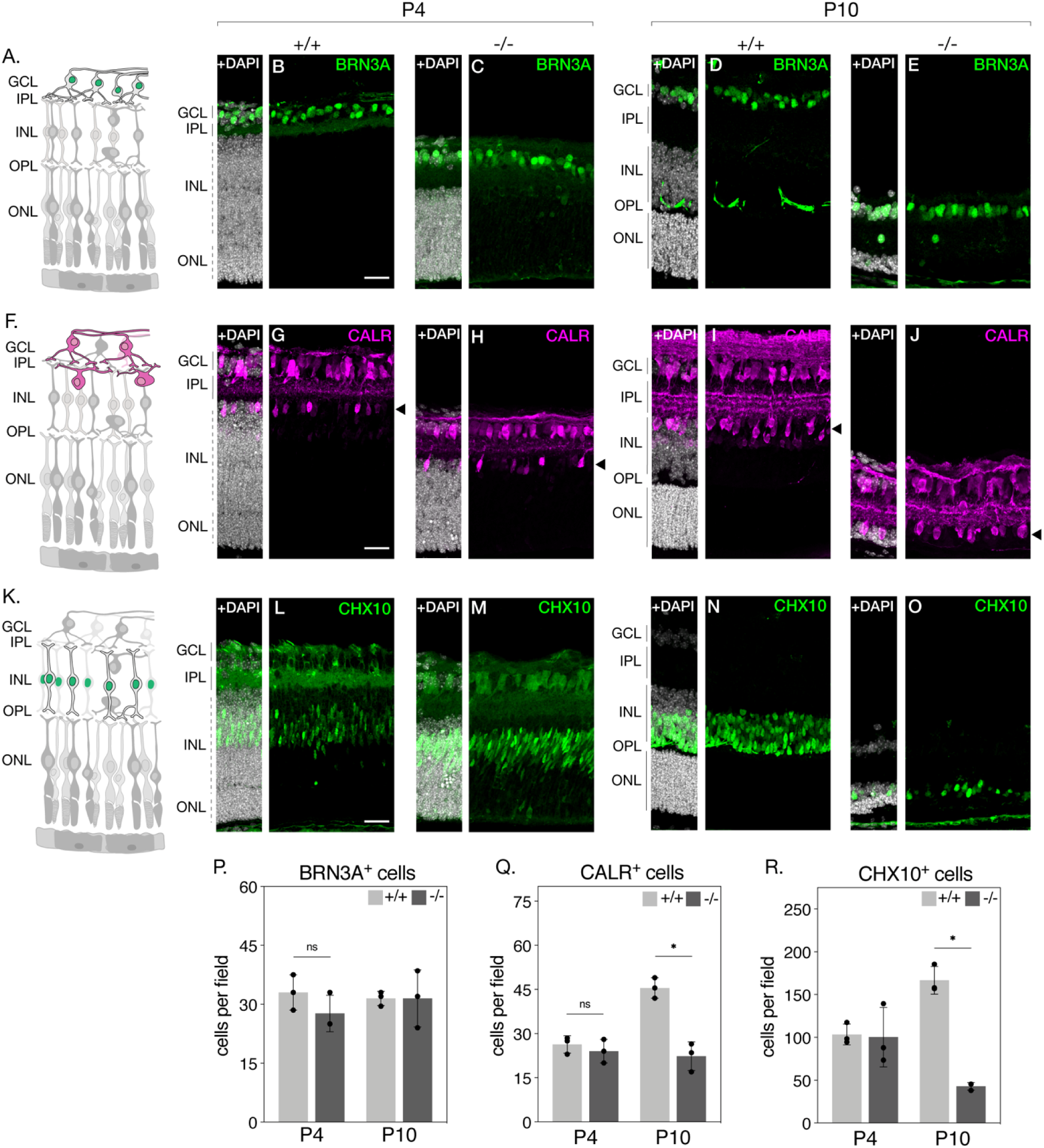
NMNAT1 loss affects survival of retinal bipolar and amacrine cells. Representative retinal sections from knockout (-/-) and floxed littermate control (+/+) mice at the indicated ages labelled with antibodies against BRN3A (**B-E**), Calretinin (CALR) (**G-J**), and CHX10 (**L-O**). Schematics in (**A)**, (**F)**, and (**K)** depict retinal neuron subtypes labelled by each respective antibody. Quantification of BRN3A- (**P**), CALR- (**Q**), and CHX10-positive cells (**R**) are shown. In (**Q)**, only CALR-positive cells on the outer side of the IPL (layer indicated by arrowheads) were counted. Data is represented as mean ± SD. n=3 biological replicates for all panels; significance determined using Student’s t-test. Scale bars, 30 μm.

### Loss of NMNAT1 impairs photoreceptor gene expression and severely perturbs early postnatal photoreceptor survival

Turning our attention to photoreceptors, we repeated the above approach with antibodies against the photoreceptor markers recoverin (anti-RCVRN) and rhodopsin (anti-RHO). While anti-RCVRN cleanly labels developing photoreceptor somas in P4 and P10 control retinas (Figure 3B, **Figure 3—Supplement 1, panel D**), we observe a complete lack of recoverin expression in knockout retinas at both ages (Figure 3C**, Figure 3—Supplement 1, panel E).** Barring a small amount of non-specific staining likely originating from the secondary antibody (**Figure 3— Supplement 1, panels A, B**), rhodopsin expression at P4 and P10 showed an identical trend to that of recoverin (Figure 3D, E**, Figure 3—Supplement 1, panels H,I**).

**Figure 3.**
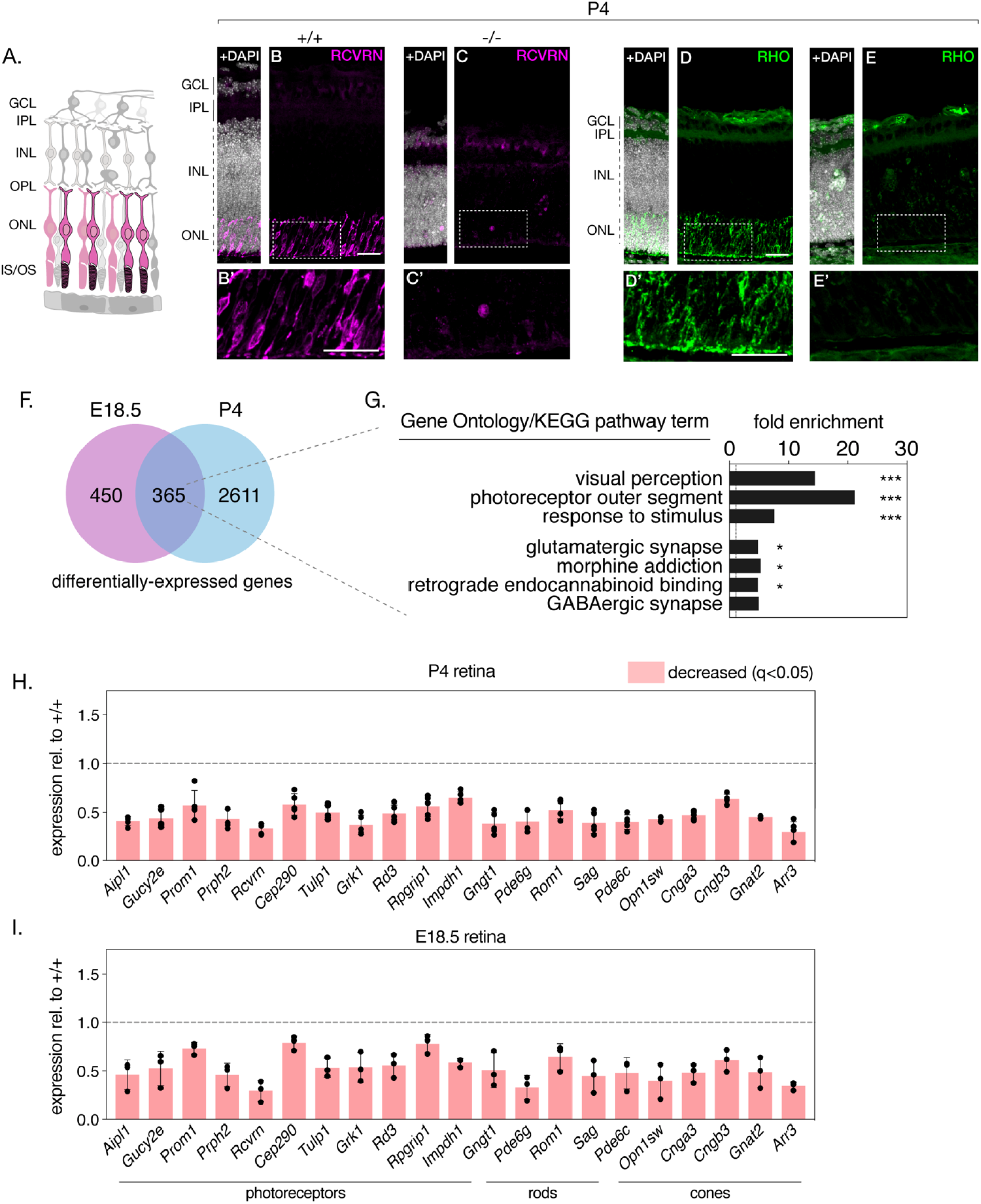
NMNAT1 loss severely affects early postnatal photoreceptor gene expression and survival. Representative retinal sections from knockout (-/-) and floxed littermate control (+/+) mice at the indicated ages labelled with antibodies against recoverin (RCVRN) (**B-C**) or rhodopsin (RHO) (**D-E**). Schematic in (**A)** depicts retinal neuron subtypes labelled by the recoverin and rhodopsin antibodies. (**F**) Comparison of differentially-expressed genes in E18.5 and P4 knockout retinas as assessed by RNA-sequencing. (**G**) GSEA of differentially-expressed genes present in E18.5 and P4 knockout retinas. (**H, I**) Relative expression of indicated genes in P4 and E18.5 knockout retinas as assessed by RNA-sequencing. n=3 biological replicates for (**B-E**), n=5 biological replicates for (**H**), n=3 biological replicates for (**I**). Corresponding zoom panels are indicated with dotted rectangles. Scale bars, 30 μm.

Intrigued by the magnitude of recoverin and rhodopsin loss and hypothesizing defects in the expression of other retinal proteins in our knockout, we comprehensively profiled the transcriptomes of knockout and control retinas at two timepoints— pre-degeneration (E18.5) and during degeneration (P4) and using RNA-sequencing. At P4, this analysis reveals 2,976 differentially-expressed genes in NMNAT1 knockout retinas (**Figure 3—Supplement 2, panel B**), several of which we validated using RT-qPCR (**Figure 3—Supplement 1, panel C**). Consistent with the lack of recoverin and rhodopsin staining at this age, gene set enrichment analysis (GSEA) of P4 differentially-expressed genes (DEGs) reveals several large, highly-overrepresented clusters of downregulated photoreceptor-related genes including both recoverin and rhodopsin (**Figure 3—Supplement 2, panel C**). Strikingly, among 815 DEGs in E18.5 knockout retinas, a similar cluster of downregulated genes associated with visual perception and the photoreceptor outer segment was observed (**Figure 3—Supplement 3, panels B, C**). Combining both RNA-sequencing datasets reveals a group of 365 DEGs in knockout retinas common to both timepoints (Figure 3F). Importantly, GSEA on this gene set reveals highly-overrepresented clusters of photoreceptor and synapse associated genes (Figure 3G), and further analysis identifies a core set of 21 photoreceptor-associated genes which are significantly downregulated in E18.5 and P4 NMNAT1 knockout retinas (Figure 3H, I). Notably, this set includes rod-specific (e.g. *Gngt1*), cone-specific (e.g. *Opn1sw*, *Cnga3*) and photoreceptor-specific (e.g. *Prph2*, *Rcvrn*, *Aipl1*) genes of diverse function, many of which have important roles in photoreceptor development and function. Consistent with a specific transcriptional effect on photoreceptors, we confirmed that expression of several well-known ganglion cell, amacrine/horizontal cell, and bipolar cell specific genes was largely unchanged in NMNAT1 knockout retinas at either tested age (**Figure 3—Supplement 4**). Altogether, these results implicate NMNAT1 in gene regulation during late-stage retinal development, suggests photoreceptor-specific transcriptional dysregulation as a driver of the severe photoreceptor phenotype in NMNAT1 deficient retinas.

Beyond affecting photoreceptor-specific gene expression, we also note downregulation of 6 synapse-associated genes (*Stx3*, *Syngr1*, *Cln3*, *Scamp5*, and *Sv2b*) in both E18.5 and P4 knockout retinas (**Figure 3—Supplement 5, panels C, D**), consistent with disruptions to outer plexiform layer formation in P4 knockout retinas on histology and on staining with the synapse marker synaptophysin (anti-SYPH) (**Figure 3—Supplement 5**).

### Loss of NMNAT1 during retinal development triggers multiple cell death pathways

As NMNAT1 deficiency drastically impairs the postnatal survival of photoreceptor, bipolar, and amacrine retinal neurons, we sought to determine the mechanisms by which these cells degenerate. To this aim, we began by staining retinal sections with an antibody against activated caspase-3 (AC3). While P0 knockout retinas show little AC3 staining compared to controls— consistent with grossly normal retinal morphology at this age—P4 knockout retinas show robust AC3 immunoreactivity in the inner and outer nuclear layers (Figure 4A-D). As expected, most AC3-immunoreactive (AC3^+^) cells display nuclear chromatin condensation (‘pyknosis’) characteristic of dying cells; however, staining also reveals a population of pyknotic nuclei not immunoreactive to AC3 (AC3^-^) (Figure 4A’-D’**, arrows**). Interestingly, these pyknotic, AC3^-^ nuclei were sparsely present in P0 and P4 control retinas (Figure 4A,C) and to a larger extent in P0 and P4 knockout retinas (Figure 4B,D). Quantification (Figure 4E) reveals a general trend of cell death consistent with our histology and cell-marker investigations: nuclear pyknosis is slightly elevated in P0 knockout retinas compared to controls, peaks at P4 where we observe robust retinal degeneration, and is virtually absent by P10, by which time the majority of outer and inner nuclei in the knockout are lost (Figure 1D). Interestingly, we observe roughly equal amounts of AC3^+^ and AC3^-^ pyknotic cells in P0 knockout retinas, whereas by P4 AC3^-^ pyknotic cells constitute ∼30% of pyknotic cells in knockout retinas (Figure 4E). In addition to being present at all tested ages and following the same general trend as AC3^+^ pyknotic cells, AC3^-^ pyknotic cells often appear in distinct clusters (Figure 4D’), distinguishable from the more evenly dispersed AC3^+^ pyknotic cells. These results suggest the activation of at least two distinct cell death pathways in NMNAT1 knockout retinas between P0 and P4.

**Figure 4.**
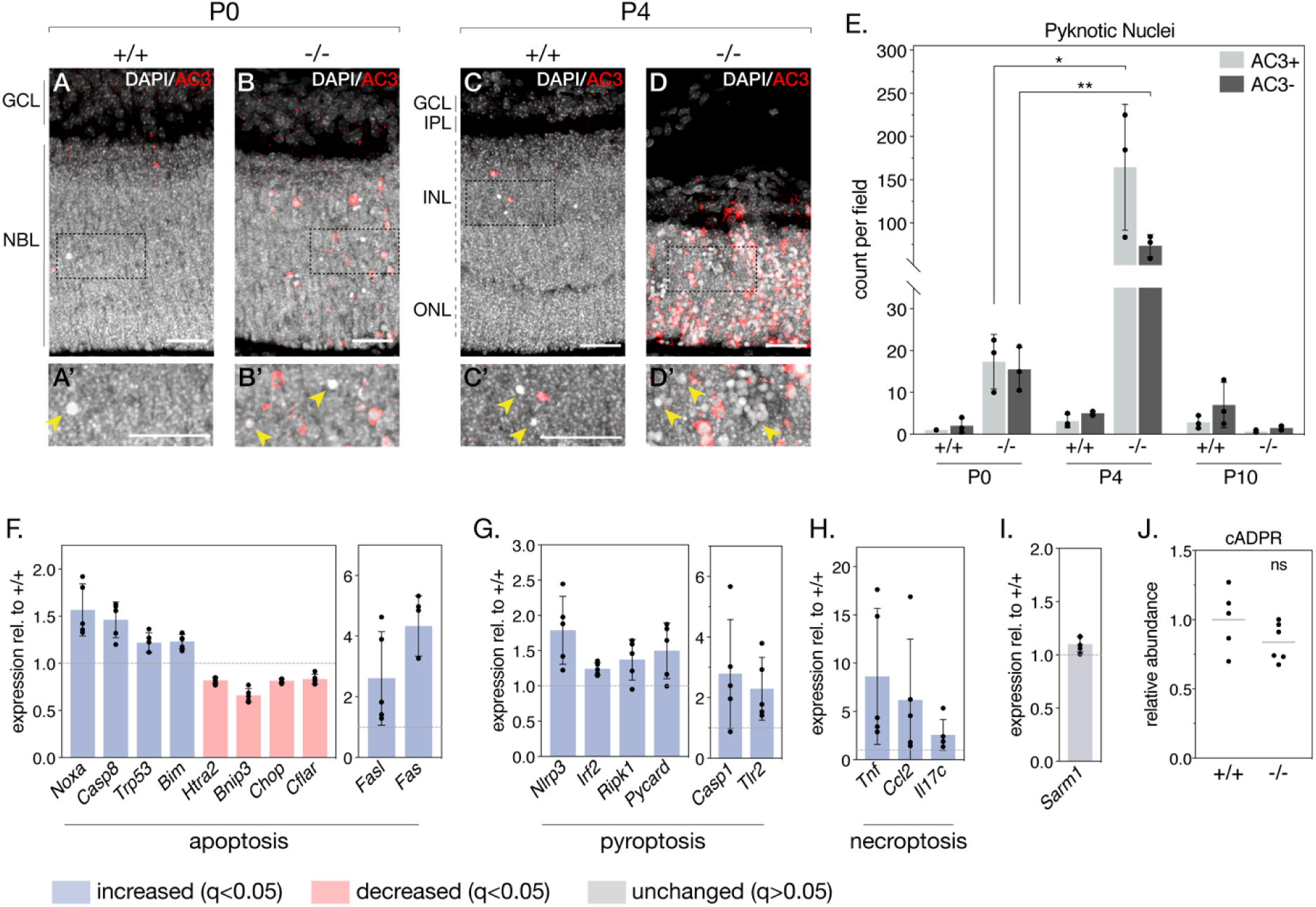
NMNAT1 loss causes activation of multiple cell death pathways in the retina. **(A-D)** Representative retinal sections from knockout (-/-) and floxed littermate control (+/+) mice at the indicated ages labelled with an antibody against active Caspase-3 (AC3). Corresponding zoom panels are indicated with dotted rectangles. Arrows denote pyknotic, AC3-negative nuclei. (**E)** Quantification of pyknotic nuclei in sections from knockout and control mice at the indicated ages, grouped by presence (AC3+) or absence (AC3-) of active Caspase-3 labeling. Relative expression of several apoptotic (**F**), pyroptotic (**G**), and necroptotic (**H**) genes in P4 knockout retinas as assessed by RNA-sequencing. (**I)** Relative expression of *Sarm1* in P4 knockout and control retinas as assessed by RNA-sequencing. (**J)** Relative abundance of cyclic-ADP-ribose (cADPR) in P4 knockout and control retinas as measured by mass spectrometry (grey bars represent means). Data are represented as mean ± SD. Significance determined using Tukey’s multiple comparisons test for (**E**), DESeq2 for (**F-I**) (see methods), or Student’s t-test for (**J**). n=3 biological replicates per condition for (**A-E**), n=5 biological replicates for (**F-I**), n=6 biological replicates (one outlier removed) for (**J**). Scale bars, 30 μm.

To more comprehensively characterize NMNAT1-associated cell death and identify possible caspase 3-independent cell death pathways in our knockout, we leveraged the E18.5 and P4 RNA-sequencing datasets mentioned above. This allowed us to systematically assay the expression of a collection of genes associated with several major cell death pathways (**Figure 4—Supplement 2**). Consistent with AC3 staining, we observe deregulation of a collection of apoptosis-related genes in P4 knockout retina, including significant increases in *Noxa* and *Fas*, two pro-apoptotic genes previously associated with cell death in NMNAT1-deficient retinas (Kuribayashi et al., 2018) (Figure 4F). Notably, two of these genes—*Noxa* and *Chop*—are also significantly deregulated at E18.5, prior to significant retinal degeneration (**Figure 4— Supplement 1, panel A**).

In addition to transcriptional signatures of apoptosis, we identified upregulation of a collection of genes associated with pyroptosis in P4 NMNAT1 knockout retinas (Figure 4G). Pyroptosis is characterized by assembly of a multi-protein complex called the ‘inflammasome,’ which ultimately cleaves and activates the pore-forming members of the gasdermin family of proteins to elicit lytic cell death in response to a variety of perturbations (McKenzie et al., 2020). Interestingly, we find upregulation of all three classical inflammasome components—*Nlrp3*, *Casp1*, and *Pycard* (ASC)—in P4 knockout retinas (Figure 4G), with *Nlrp3* upregulation at E18.5 as well (**Figure 4—supplement 1, panel B**). In addition, we observe significant increases in *Irf2*, a transcriptional activator of gasdermin D (Kayagaki et al., 2019), as well as pyroptosis-associated proteins *Ripk1* and *Tlr2* at P4 (Figure 4G). Notably, expression of *Tlr2* and related protein *Tlr4* is significantly elevated in E18.5 knockout retinas (**Figure 4—Supplement 1, panel B**). Finally, we also observed dysregulation of several genes associated with necroptosis (Figure 4H) and ferroptosis (**Figure 4—Supplement 2, panel D**) in P4 knockout retinas; while none of these genes were significantly upregulated at E18.5, we do observe an early induction of necroptosis-associated protein *Nox2* at this age (**Figure 4—Supplement 1, panel C**).

Recently, photoreceptor cell death in a postnatally-induced global NMNAT1 knockout mouse was shown to depend heavily on the activity of the pro-degenerative axonal protein SARM1 (Sasaki et al., 2020a). Reasoning SARM1 as the culprit behind the caspase 3-independent cell death in our model, we checked *Sarm1* expression in our RNA-seq data and assayed SARM1 activity by measuring levels of its catalytic product cyclic ADP-ribose (cADPR) using targeted mass spectrometry in P4 and E18.5 NMNAT1 knockout and control retinas. Surprisingly, we found no significant changes in SARM1 expression (Figure 4I**, Figure 4—Supplement 1, panel D**) or activity (Figure 4J**, Figure 4—Supplement 1, panel E**) at either tested age. Overall, these data reveal that activation of multiple cell death pathways underlies the early and severe degeneration observed in NMNAT1 knockout retinas, and suggest pyroptosis and apoptosis as drivers of this degeneration.

### Global metabolic alterations in NMNAT1 deficient retinas

To identify possible mechanisms for the severe and cell type-specific retinal degeneration in our model, we next sought to characterize global metabolic consequences of embryonic NMNAT1 deletion in the retina. To this end, we used targeted liquid chromatography-tandem mass spectrometry (LC-MS/MS) to quantify levels of ∼112 cellular metabolites spanning many essential biochemical pathways in NMNAT1 knockout and control retinas at pre- and post-degenerative timepoints matching that of our RNA-sequencing analyses (E18.5 and P4). While LC-MS/MS analysis reveals no significant changes in E18.5 knockout retinas compared to controls, analysis at P4 reveals significantly altered levels of 39 metabolites in knockout retinas (Figure 5**, Figure 5—Supplement 1**). Metabolite set enrichment analysis (MSEA) identifies potential disruption of several diverse biochemical pathways including amino acid metabolism, glycolysis/gluconeogenesis, nicotinate and nicotinamide metabolism, and purine metabolism (Figure 5B).

**Figure 5.**
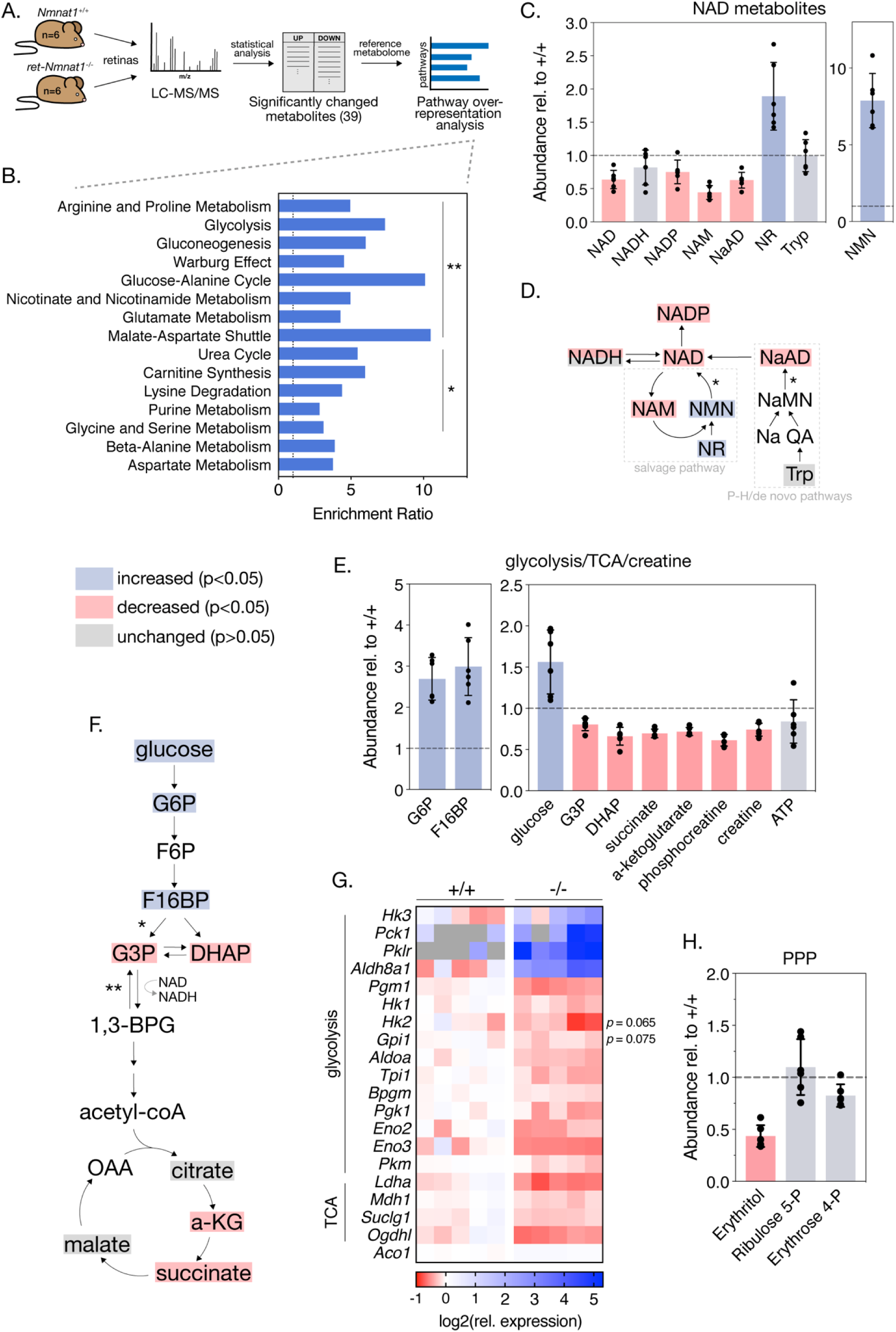
Loss of NMNAT1 impairs retinal central carbon metabolism. (**A**) Schematic illustrating metabolomics experimental approach. (**B**) Results of metabolite set enrichment analysis (MSEA) on significantly changed metabolites in P4 knockout retinas. (**C**) Relative abundance of NAD^+^ pathway metabolites in P4 knockout retinas as assessed by mass spectrometry. (**D**) Schematic illustrating the major mammalian NAD^+^ synthesis pathways colored according to metabolite changes in (**C**); NMNAT1-catalyzed steps are indicated with asterisks. (**E**) Relative abundance of glycolysis, TCA cycle, and creatine metabolites in P4 knockout retinas as assessed by mass spectrometry. (**F**) Schematic depicting abbreviated glycolysis/TCA cycle pathway colored according to metabolite changes in (**E**); aldolase-catalyzed step is indicated by a single asterisk, while GAPDH-catalyzed step is marked with a double asterisk. (**G**) Heatmap of log-transformed relative expression of a set of glycolysis/TCA cycle genes in P4 knockout (-/-) and control retinas (+/+) as assessed by RNA-sequencing. (**H**) Relative abundance of pentose-phosphate pathway (PPP) metabolites in P4 knockout retinas as assessed by mass spectrometry. Data are represented as mean ± SD. n=6 biological replicates for (**B,C,E,H**), n=5 biological replicates for (**G**).

NMNAT1 knockout retinas show specific metabolic disruptions to NAD^+^ biosynthesis pathways. At P4, knockouts show a ∼40% reduction of total retinal NAD^+^ levels and levels of nicotinic acid adenine dinucleotide (NaAD), the other catalytic product of NMNAT1 (Figure 5C, D). Levels of the downstream metabolites NADP and nicotinamide (NAM) were decreased by ∼25% and ∼55%, respectively, while levels of NADH were slightly decreased (p > 0.05) (Figure 5C). As expected, we observe significant accumulation of NAD precursors nicotinamide riboside (NR) and nicotinamide mononucleotide (NMN) in P4 knockout retinas (Figure 5C); however, we observe no significant changes in levels of tryptophan, the starting point for de novo NAD synthesis, at this age (Figure 5C, D).

### Retinal NMNAT1 loss causes disruption of central carbon metabolism

Interestingly, the set of significantly altered metabolites in P4 knockout retinas is enriched for metabolites associated with glycolysis, gluconeogenesis, and the Warburg effect (Figure 5B). Closer examination of these pathways reveals large relative increases in levels of the upstream glycolytic metabolites glucose, glucose 6-phosphate (G6P) and fructose 1,6-bisphosphate (F16BP), as well as significant decreases in levels of dihydroxyacetone phosphate (DHAP) and glucose 3-phosphate (G3P) (Figure 5E, F), strongly suggesting a disruption to glucose utilization. Consistent with such an effect, levels of the TCA cycle intermediates alpha-ketoglutarate (a-KG) and succinate are decreased by ∼30% in P4 knockout retinas (Figure 5E, F). Further in line with disruptions to downstream mitochondrial metabolism, we observe reduction of several acylcarnitine species at this age as well (**Figure 5—Supplement 1, panel A**). Although we observe decreased levels of the ATP-recycling metabolites phosphocreatine and creatine at P4, retinal ATP levels at this age are slightly but not significantly reduced (p = 0.5) (Figure 5F).

To further investigate possible glycolytic disruptions in P4 knockout retinas, we assayed the expression of a collection of glycolysis and TCA cycle enzymes in P4 knockout and control retinas using our RNA-sequencing dataset. Indeed, this analysis reveals broad transcriptional changes in 16 glycolytic and 4 TCA cycle enzymes in P4 knockout retinas (Figure 5G). Notably, two of these changes—upregulation of the aldehyde dehydrogenase *Aldh8a1* and downregulation of the TCA cycle enzyme *Ogdhl*—are also present in RNA-sequencing results from the pre-degenerative (E18.5) timepoint (**Figure 5—Supplement 1, panel B**). Finally, consistent with disruption of glycolytic flux and reduced NADP levels in knockout retinas, mass spectrometry suggests possible disruption of the pentose phosphate pathway (PPP) evidenced by decreased levels of erythritol at this age (Figure 5H).

### NMNAT1 loss disrupts retinal purine metabolism and a subset of amino acids

As MSEA also indicated possible disruptions of purine metabolism and several amino acid pathways in P4 NMNAT1 knockout retinas (Figure 5B), we closely examined levels of a collection of metabolites representing most major nucleotide and amino acid metabolites in P4 knockout and control retinas (**Figure 5—Supplement 2**). Interestingly, while levels of the pyrimidine nucleotide derivatives uracil, UDP, and cytidine were unchanged at this age, levels of the purine nucleotide precursor xanthine, guanine, and GMP were increased by ∼150%, ∼147%, and ∼58%, respectively (**Figure 5—Supplement 2, panel A**). The sole affected pyrimidine nucleotide was cytosine, which showed an impressive ∼352% increase in P4 knockout retinas as compared to controls (**Figure 5—Supplement 2, panel A**). In addition to defects in purine metabolism, we observe significant changes in a subset of 10 amino acids and amino acid derivatives in P4 knockout retinas including acetyl-asparagine and acetyl-lysine (**Figure 5— Supplement 2, panel B**).

Overall, these metabolomics results suggest specific disruptions to central carbon, purine nucleotide, and amino acid metabolism as potential causes for severe retinal degeneration in the absence of NMNAT1.

## DISCUSSION

### NMNAT1-deficiency is associated with early and severe retinal degeneration involving multiple cell types

In this study, we demonstrate that retinal NMNAT1 deficiency in mice leads to severe degeneration of photoreceptor, bipolar, and amacrine neurons soon after birth. In general, this phenotype is consistent with several recent studies reporting partial or complete ablation of retinal NMNAT1, which report retinal degeneration beginning within the first postnatal week and largely complete by one month of age (Wang, et al., 2017; Eblimit et al., 2018). While the present study describes a relatively rapid timeline of NMNAT1-associated retinal degeneration, it furthers the characterization of this phenotype in two important ways: first, by assessing the survival of specific retinal neuron subtypes, and second, by systematically examining retinal cell death pathways triggered by NMNAT1 deletion.

Examination of specific cell types in NMNAT1 knockout retinas reveals that, while photoreceptors are likely the primary targets of NMNAT1-associated pathology, retinal bipolar and amacrine cells are also sequentially and significantly affected by loss of NMNAT1 during retinal development, while ganglion cells appear unaffected until later stages. Interestingly, this differential sensitivity is mirrored in our transcriptomics analyses, which demonstrate robust deregulation of a cluster of photoreceptor and several bipolar cell-specific genes but relatively few changes in amacrine or ganglion specific genes. These results stand partially in contrast to a recent study reporting *ex vivo* knockdown of *Nmnat1* in retinal explant cultures, which reported thinner INLs but no changes in numbers of HuC/D-expressing amacrine cells or PKCa-expressing bipolar cells in NMNAT1-deficient explants (Kuribayashi et al., 2018). However, this study did report reduced numbers of PNR-positive photoreceptor cells in NMNAT1-deficient explants, and we believe that differences regarding amacrine and bipolar cell effects can be explained by the fact that *Nmnat1* expression in explants was knocked down relatively late in development (E17.5) as compared to the E9.5 activation of *Six3-Cre* (Furuta et al, 2000).

While we were unable to successfully determine retinal NMNAT1 distribution using our polyclonal NMNAT1 antibody, previously published results showing RT-qPCR of *Nmnat* levels in flow-sorted rod photoreceptors suggest that NMNAT1 is the predominantly expressed NMNAT isoform in rod photoreceptors (Kuribayashi et al., 2018). Combined with our data confirming a lack of transcriptional upregulation of either *Nmnat2* or *Nmnat3* in NMNAT1 knockout retinas (**Figure 3—Supplement 1**), one possible explanation for the cell-type specific degeneration we observe is that relatively higher levels of NMNAT2/3 in inner retinal neurons can partially compensate for the NAD^+^ deficit caused by loss of NMNAT1. Overall, our results demonstrate that NMNAT1 is crucial for the early survival of photoreceptors, bipolar cells, and amacrine cells and suggest retinal cell-type specific requirements of NAD^+^ metabolism.

### NMNAT1-deficient retinas activate multiple cell death pathways

Previous reports of cell death in NMNAT1-deficient retinas center on two potentially contrasting mechanisms: the aforementioned *ex vivo* study outlined a role for *Noxa* and *Fas*-associated, caspase-3 dependent apoptosis in NMNAT1-deficient explants (Kuribayashi et al., 2018), while a recent study found the death of mature photoreceptors after global NMNAT1 deletion to be solely dependent on the neuronal NADase SARM1 (Sasaki et al., 2020a). Using histological and comprehensive transcriptomic approaches, we demonstrate involvement of caspase-3 associated apoptosis in NMNAT1 knockout retinas characterized by an early and sustained upregulation of *Noxa* and deregulation of several other apoptosis-pathway genes. We extend these results by showing histological evidence of a distinct, caspase-3 independent cell death pathway which constitutes a significant portion of observed cell death and closely follows caspase-3 dependent apoptosis throughout the timepoints tested in our model.

Interestingly, using a recently validated metabolic marker of SARM1 activity (cADPR) (Sasaki et al., 2020b), we do not detect SARM1 involvement at pre- or post-degenerative timepoints in our model. That we do not find evidence of SARM1 involvement, even in the presence of increased levels of its potent activator NMN (Zhao et al., 2019; Figley et al., 2021) is unexpected but not implausible, especially considering the fact that the report implicating SARM1 in NMNAT1-associated degeneration deleted NMNAT1 in mature mice (Sasaki et al., 2020a).

Indeed, many apoptosis effectors (*Casp3*, *Casp9*, *Apaf1*, *Bcl* family members) active in the developing mammalian retina are subsequently downregulated in mature retinas, necessitating alternative death pathways for handling pathological insults at these ages (Donovan and Cotter 2002; Doonan et al., 2003; Donovan et al., 2006). Considering these results, it is wholly possible that in mature retinas with insufficient expression of necessary apoptotic and/or pyroptotic machinery, NMNAT1-associated retinal degeneration proceeds through SARM1—in such a case, it is important to note that model systems with embryonic or germline deletion or mutation of NMNAT1 are typically more representative of patients with disease-linked mutations in NMNAT1. On the other hand, it cannot be conclusively ruled out that excess cADPR produced by an active SARM1 in our model is rapidly metabolized, and recently described links between SARM1 and both pyroptosis and apoptosis (Mukherjee et al., 2015; Carty et al., 2019) leave open the possibility of cooperation between these cell death pathways in NMNAT1 knockout retinas.

Our transcriptomics results indicate dysregulation of several pyroptosis and necroptosis-related genes, notably including the entire “canonical inflammasome” (*Casp1*, *Nlrp3*, and *Pycard*), as well as the toll-like receptors *Tlr2* and *Tlr4*, at both timepoints (Figure 4). The distinct presence of nuclear pyknosis in AC3^—^ dying cells—which is generally incompatible with necroptosis but documented in pyroptotic cells (Vandenabeele et al., 2010; Murakami et al., 2012; Miao et al., 2011)—lends support to pyroptosis as a significant driver of cell death in NMNAT1-deficient retinas. Intriguingly, we do not detect proteolytic cleavage of gasdermin D—a common marker of pyroptosis—in P4 NMNAT1 knockout retinas (**Figure 4—Supplement 1**). However, recent results indicate that NLRP3 is, under certain circumstances, capable of being activated independently of gasdermin D (Gutierrez et al., 2017). Considering a recent report suggesting involvement of the PARP1-associated ‘parthanatos’ cell death pathway in NMNAT1 mutant retinas (Greenwald et al., 2021), we did not detect accumulation of poly ADP-ribose (PAR) in AC3^—^ pyknotic nuclei (data not shown), arguing against involvement of this pathway under these conditions.

In sum, we show that retinal NMNAT1 loss activates multiple cell death pathways, which contextualizes the degeneration of multiple cells types in our model and may explain the severity of NMNAT1-associated retinal degeneration in this study and others. Although the model presented here differs in important ways from NMNAT1-mutant LCA animal models, it does recapitulate some aspects of NMNAT1-linked LCA including particularly severe central retinal defects (Kumaran et al., 2017). As recent studies have discovered noncoding mutations, copy number variations, and exon duplications in NMNAT1 causing severe reduction of NMNAT1 expression in patients with ocular and extra-ocular pathologies (Coppieters et al., 2015; Bedoni et al., 2020), understanding the mechanisms of retinal degeneration in an NMNAT1 knockout model is of potential clinical significance.

### Retinal NMNAT1 loss causes diverse metabolic disruptions

Several recent studies examine levels of select metabolites in mature NMNAT1-deficient retinas (Sasaki et al., 2020a), or more broadly examine tissue metabolomes after perturbation of the NMN-synthesizing enzyme NAMPT (Lin et al., 2016; Oakey et al., 2019; Lundt et al., 2021). Our metabolomics results approximate that NMNAT1 synthesizes ∼40% of the total retinal NAD^+^ pool, which is generally consistent with a previous model (Sasaki et al., 2020a). In addition, our results strongly suggest that—via specific disruptions to central carbon, nucleotide, and amino acid metabolism—depletion of NAD^+^ synthesized in the nucleus disrupts multiple non-nuclear metabolic pathways in the retina. Whether these observations reflect a direct export of nuclear NAD^+^ to cytosolic and mitochondrial retinal compartments versus an indirect effect on cytosolic and mitochondrial cellular processes (by way of NAD^+^-dependent gene regulation, for instance) is a topic for further study.

Impaired glycolytic flux appears to be a more general feature of tissue NAD^+^ depletion, as studies reporting NAMPT inhibition or deletion in projection neurons and skeletal muscle myotubes report accumulation of glycolytic metabolites upstream of GAPDH (Oakey et al., 2019; Lundt et al., 2021). Interestingly, Lundt et al. present evidence of reversed glycolytic flux in NAMPT-inhibited myotubes, an effect which we believe may explain decreased levels of G3P and DHAP in our model. Notably, unbiased LC-MS/MS and GC-MS analyses of rod-photoreceptor-specific NAMPT knockout retinas showed signatures of mitochondrial metabolic defects—which we observe in our model as well—but detected limited evidence of glycolytic impairment (Lin et al., 2016). This suggests that retinal NMNAT1 and NAMPT depletion, despite both lowering total retinal NAD^+^ levels, produce distinct metabolic phenotypes.

Retinal neurons—and retinal photoreceptors in particular—are relatively unique in their dependence on aerobic glycolysis (the “Warburg Effect”) during both proliferative and differentiated states (Agathocleous et al., 2012; Ng et al., 2015; Chinchore et al., 2017). Our finding that retinal NMNAT1 loss is detrimental to glycolysis thus offers a potential explanation for the cell type-specific degeneration which we observe: in particular, previous results indicating that differentiation induces an increased reliance on mitochondrial OXPHOS relative to glycolysis in the retina (Agathocleous et al., 2012) might explain why ganglion and amacrine cells—which differentiate relatively early—are less sensitive to NMNAT1 loss in our model than the later born bipolar and photoreceptor cells. Indeed, even in mature retinas, glycolytic perturbations specifically affect photoreceptor health and survival (Chinchore et al., 2015; Zhang et al., 2020; Sinha et al., 2021). Furthermore, recent results linking glycolytic impairment to NLRP3 activation (Sanman et al., 2016) provide a potential explanation of non-apoptotic cell death which we observe in NMNAT1 knockout retinas.

In addition to glycolytic impairment, we detect specific defects in purine nucleotide and amino acid metabolic pathways in NMNAT1 knockout retinas. As a particularly proliferative tissue, the retina is thought to be highly reliant on adequate nucleotide and amino acid pools to support transcription and translation of cell-specific machinery (Etingof 2001; Ng et al., 2015). Some of the metabolic changes which we observe—for instance, accumulation of the purine precursor xanthine and the amino acid aspartate—appear to be more widely associated with NAD^+^ insufficiency or retinal degeneration (Du et al., 2014; Lin et al., 2016; Oakey et al., 2019). On the other hand, we also identify a collection of metabolic changes in these pathways which are not reported in NAMPT-deficient retinas (Lin et al., 2016), potentially explaining differences in retinal phenotypes between these two models, to be explored more in future studies.

### A potential role for NMNAT1 in photoreceptor terminal differentiation

The tightly coordinated and stereotypical differentiation of retinal neuron subtypes from a common progenitor pool has been extensively studied—complementing classical birth-dating studies, recent investigations have begun to explore the massive epigenetic regulation necessary for the development of the mammalian retina (Swaroop et al., 2010; Aldiri et al., 2017; Raeisossadati et al., 2021). One of the most surprising findings of the present study is an early and sustained transcriptional downregulation of a subset of photoreceptor- and synapse-specific genes in NMNAT1 knockout retinas. Beyond these transcriptional disruptions, we show near complete absence of rhodopsin and recoverin protein in P4 NMNAT1 knockout retinas. While NMNAT1 has previously been implicated in gene regulation through direct interaction with SIRT1 and PARP1 at gene promoters (Zhang et al., 2012; Song et al., 2013) and NMNAT1 knockdown was shown to influence apoptotic gene expression by potentially modulating histone acetylation (Kuribayashi et al., 2018), no *in vivo* role for NMNAT1 in retinal developmental gene regulation has yet been described.

Photoreceptors are among the last retinal cell types to fully develop and are generated in two broad phases: an early cell-fate commitment mediated by several well-characterized transcription factors including OTX2, NRL, and CRX, and a later phase (“terminal differentiation”) comprising expression of a host of specialized phototransduction genes and growth of light-sensing cellular structures (Swaroop et al., 2010; Brzezinski and Reh, 2015; Daum et al., 2017). Interestingly, we do not observe transcriptional changes in OTX2, NRL, CRX, or related genes in NMNAT1 knockout retinas at either tested age. This fact, combined with grossly normal retinal morphology at P0, suggests that NMNAT1 is required for terminal differentiation but not early specification or proliferation of retinal photoreceptor cells. Increased abundance of acetyl-lysine in knockout retinas on mass spectrometry (**Figure 6B**) and GO enrichment of several genes associated with DNA methylation, epigenetic regulation, and chromatin silencing in P4 knockout retinas (**Figure 3—Supplement 2**) support a potential role for NMNAT1 in the epigenetic regulation of photoreceptor terminal differentiation, a phenomenon recently shown to feature genome-wide methylation and acetylation events (Aldiri et al., 2017).

In conclusion, this study presents the most comprehensive evaluation of NMNAT1-associated retinal dysfunction to date and suggests crucial roles for nuclear NAD^+^ in the proper development and early survival of the mammalian retina. While we provide evidence that the early and severe retinal degeneration associated with NMNAT1 loss involves multiple cell types and death pathways, it appears that this severe phenotype stems from two major problems: 1.) metabolic defects likely caused by insufficient NAD^+^ for retinal proliferative metabolism, and 2.) gene regulation defects potentially caused by insufficient nuclear NAD^+^ in developing photoreceptors. Considering links between metabolic state and differentiation in the retina and recently discovered roles of compartmentalized NAD^+^ in non-retinal cell differentiation (Agathocleous et al., 2012; Agathocleuous et al., 2013; Ryu et al., 2018), these two problems may not be mutually exclusive. Further study of NMNAT1-associated retinal dysfunction should focus on evaluating the relative contributions of metabolic and genetic deficits to the overall pathology and testing the hypothesis that NMNAT1 functions to integrate retinal energy metabolism and gene regulation.

## MATERIALS AND METHODS

### Animal Model Generation, Husbandry, and Genotyping

*Nmnat1*^fl/fl^ mice were described previously (Conforti 2011) were thoroughly backcrossed with wild-type 129/SV-E mice (Charles River Laboratories, Wilmington, MA) prior to analyses. Conditional knockout mice were generated by crossing *Nmnat1*^fl/fl^ mice with transgenic mice expressing Cre recombinase under a *Six3* promoter (*Six3-Cre*) (Christiansen et al., 2011). Crosses yielded heterozygous *Nmnat*^fl/wt^ and *Six3-Cre Nmnat*^fl/wt^ offspring, which were further crossed with *Nmnat1*^fl/fl^ mice to yield conditional knockout (*Six3-Cre Nmnat*^fl/fl^) and littermate control (*Nmnat1*^fl/fl^) mice at approximately Mendelian ratios. Experimental animals were periodically backcrossed with wild-type 129/SV-E mice to maintain genetic integrity. Animals were maintained under standard 12 hour light/dark cycles with food and water provided *ad libitum*. All experimental procedures involving animals were approved by the Institutional Animal Care and Use Committee (IACUC) of West Virginia University.

Animals were genotyped using polymerase chain reaction (PCR) of genomic DNA from ear punch biopsies. Primer sequences for detection of *Six3-Cre* transgene and *Nmnat1* 5’ and 3’ loxP sites are listed in **Supplementary Table 1**, and primers were added to the PCR reaction at a final concentration of 0.75 µM. The thermocycling conditions were 95°C for 2 minutes, 35 cycles of 95°C for 30s, 58°C for 30s, 72°C for 45s, and a final extension step of 72°C for 5 minutes.

### NMNAT1 Antibody Generation

A rabbit polyclonal antibody against amino acids 111-133 of mouse NMNAT1 (sequence CSYPQSSPALEKPGRKRKWADQK) was generated by Pacific Immunology Corp. (Pacific Immunology, Ramona, CA). The affinity-purified antibody was confirmed to recognize NMNAT1 in cell culture and retinal lysate (**Figure 1—Supplement 1**), and did not recognize overexpressed FLAG-NMNAT2 (data not shown).

### Mammalian Cell Culture and Transfection

HEK293T cells were maintained in 1:1 DMEM/F-12 culture media (Thermo Fisher Scientific, Waltham, MA) in a sterile incubator at 37°C and 5% CO_2_. Transient transfection of FLAG-NMNAT1 and FLAG-NMNAT2 constructs (see Supplemental Table 2) was performed at ∼60% confluence using TransIT^®^-LT1 transfection reagent (Mirus Bio, Madison, WI) according to manufacturer’s instructions. Cells were harvested 48 hours after transfection on ice with 1X Hank’s balanced salt solution (HBSS) (Thermo Fisher Scientific) and cell pellets were processed for western blotting as described below.

### Retinal Histology and Thickness Quantification

To evaluate gross retinal histology, mice at the indicated ages were sacrificed and their eyes were gently removed into 1ml Excalibur’s Alcoholic z-fix (Excalibur Pathology Inc., Norman, OK). Subsequent fixation, paraffin embedding, sectioning, and hematoxylin and eosin (H&E) staining were performed by Excalibur Pathology. H&E-stained sections were imaged on a Nikon Eclipse Ti microscope with DS-Ri2 camera (Nikon Instruments, Melville, NY). Retinal thickness was measured using Nikon NIS-Elements software at 4 equidistant points along the outer retinal edge to either side of the optic nerve, where retinal thickness is defined as the length of a line orthogonal to the outer retinal edge and terminating at the inner retinal edge. Thickness was quantified using 6 technical replicates per animal and 3 biological replicates per genotype.

### Western Blotting

For determination of protein levels by western blot, retinas were homogenized in cold lysis buffer (1X phosphate-buffered saline pH 7.4 (Thermo Fisher Scientific), Pierce^®^ protease inhibitor, EDTA-free (Thermo), Calbiochem^®^ phosphatase inhibitor cocktail (EMD Biosciences, La Jolla, CA)) using a Microson^®^ ultrasonic cell disruptor. Protein concentration was determined using a Nanodrop ND-1000 spectrophotometer (Thermo Fisher Scientific), after which Laemmli sample buffer was added to a final concentration of 1X (2% SDS, 0.05M Tris-HCl pH 6.8, 10% glycerol, 1% β-mercaptoethanol, 2 mM dithiothreitol (DTT), 0.001% bromphenol blue) and samples were boiled for 5 minutes. Equal amounts of protein were separated on anykD® mini-PROTEAN TGX polyacrylamide gels (Bio-Rad, Hercules, CA) and transferred to an Immobilon-FL PVDF membrane (Millipore, Burlington, MA). Membranes were blocked with Odyssey^®^ blocking buffer (LI-COR Biosciences, Lincoln, NE) for 1 hour at room temperature and incubated with primary antibodies diluted in 50/50 blocking buffer/1X PBST (1X PBS + 0.1% Tween-20) at 4°C overnight on a bidirectional rotator. A list of primary antibodies, sources, and dilutions can be found in **Supplemental Table 2**. Following primary antibody incubations, membranes were washed in 1X PBST and incubated with goat anti-rabbit Alexa Fluor 680 and/or goat anti-mouse DyLight 800 secondary antibodies (Thermo Fisher Scientific) for 1 hour at room temperature. Membranes were subsequently washed in 1X PBST and imaged using an Odyssey Infrared Imaging System (LI-COR Biosciences).

### Quantitative Reverse Transcriptase-PCR (RT-qPCR)

To determine relative gene expression using RT-qPCR, retinas were collected and total RNA was isolated as described below. cDNA was synthesized with a RevertAid^®^ cDNA Synthesis Kit (Thermo Fisher Scientific) per manufacturer instructions, starting with 1µg total RNA/sample and using random hexamer primers. Primers for targets of interest were designed using NCBI Primer-BLAST (https://www.ncbi.nlm.nih.gov/tools/primer-blast/) to straddle at least one intron of length >500bp. qPCR reactions were performed using Brilliant II SYBR® Green qPCR master mix with low ROX (Agilent Technologies, Cedar Creek, TX) and monitored on a Stratagene® Mx3000P cycler (Agilent Technologies). Prior to use, primers were validated (by examining melt curves and agarose gel electrophoresis) to amplify single products of expected size. Three technical replicates per target per animal were performed, and averages from three animals per genotype are reported. All expression values were normalized to the geometric average of three housekeeping genes (*Hmbs*, *Ppia*, and *Ywhaz*).

### Immunofluorescent Staining

After euthanasia, eyes were gently removed, punctured, and immersed in 4% paraformaldehyde fixative (4% paraformaldehyde in 1X PBS) (Electron Microscopy Sciences, Hatfield, PA) for 15 minutes, after which the cornea was removed and the eye was fixed for an additional 45 minutes at room temperature with gentle agitation. Subsequently, eyes were washed in 1X PBS and incubated in a dehydration solution (20% sucrose in 1X PBS) for at least 12 hours at 4°C. After dehydration, samples were incubated in a 1:1 mixture of 20% sucrose and Tissue-Tek^®^ O.C.T. compound (Sakura Finetek, Torrance, CA) for at least 1 hour before being transferred to 100% O.C.T. and flash frozen. 16 µm sections were cut using a Leica CM1850 cryostat (Leica Biosystems, Nussloch, Germany) and mounted on Superfrost Plus slides (Fisher Scientific). For immunofluorescent staining, retinal sections were briefly rinsed with 1X PBS and incubated in blocking buffer (10% normal goat serum, 0.5% Triton X-100, 0.05% sodium azide in 1X PBS) for 1 hour at room temperature. Following blocking, sections were incubated with the indicated primary antibodies (diluted in buffer containing 5% normal goat serum, 0.5% Triton X-100, 0.05% sodium azide in 1X PBS) overnight at 4°C. The next day, sections were washed with 1X PBST and incubated with DAPI nuclear stain (Thermo Fisher Scientific), goat anti-rabbit Alexa Fluor-568 and/or goat anti-mouse Alexa Fluor-488 (Thermo Fisher Scientific) secondary antibodies for 1 hour at room temperature. For antibody information and dilutions, see **Supplemental Table 2**. Finally, sections were washed, cover-slipped with Prolong Gold^®^ antifade reagent (Thermo Fisher Scientific), and imaged on a Nikon Eclipse Ti laser scanning confocal microscope with C2 camera (Nikon Instruments). Experimental and control sections were imaged using identical laser intensity and exposure settings. All fluorescent images represent maximum intensity z-projections generated using ImageJ with the Bio-Formats plugin (https://imagej.net/Bio-Formats).

### Retinal Cell Type and Caspase-Positive Cell Quantification

To estimate numbers of pyknotic and caspase-3 positive/negative cells in retinas of knockout and control mice, pyknotic nuclei were manually counted in 212.27 x 212.27 µm^2^ regions of maximum intensity projection fluorescent images from the central retina. Once these pyknotic nuclei were identified, the number also labeled with active-Caspase-3 were manually counted. Counts from two technical replicates per animal were averaged, and averages from three animals per genotype are reported. Counts of retinal cell subtypes were estimated by manually counting marker-positive cells in 318.2 x 318.2 µm^2^ regions of central retina, averaged between two technical replicates per animal and three animals per genotype. All counts were obtained using ImageJ. For antibody information, see **Supplemental Table 2**.

### Retinal RNA Extraction, Sequencing, and Analysis

Following euthanasia and eye enucleation, retinae were quickly dissected and transferred into ultra-sterile microcentrifuge tubes containing a small amount of Trizol^®^ reagent (Thermo Fisher Scientific), and flash frozen on dry ice. Total RNA was extracted by homogenizing thawed samples with a handheld homogenizer and incubating for 5 min. at room temperature. 20µl chloroform was added to each sample, samples were briefly vortexed, incubated at room temperature for 4 min., and spun at 12,000 rpm for 10 min. at 4°C. The aqueous layers were removed to separate tubes containing 50µl isopropanol and incubated at room temperature for 10 min. with occasional agitation. Finally, samples were again spun at 12,000 rpm for 10 min. at 4°C, supernatants were removed, and pellets were washed three times with 75% ethanol, dried, and resuspended in DEPC-treated water. Whole-transcriptome sequencing was performed by Macrogen Corp. (Macrogen USA, Rockville, MD) using an Illumina TruSeq Stranded mRNA Library Prep Kit on a NovaSeq6000 S4 to a depth of 100M total reads per sample. Read quality was verified using FastQC (https://www.bioinformatics.babraham.ac.uk/projects/fastqc/) and adapters were trimmed using the Bbduk utility of the BBTools package (http://sourceforge.net/projects/bbmap/). Read alignment was performed using HISAT2 2.1.0 (Kim et al., 2015) and transcripts were assembled and quantified using StringTie 1.3.6 (Pertea et al., 2015; Pertea et al., 2016). Differentially expressed genes were identified using the DESeq2 R package (Love et al., 2014).

### Targeted Steady-State Metabolomics using LC-MS/MS

Following euthanasia and eye enucleation, retinae were quickly dissected into Hank’s Balanced Salt Solution (HBSS) and flash frozen on liquid nitrogen. Metabolites were extracted according to previously described protocols (Grenell et al., 2019; Yam et al., 2019). Metabolite extracts were analyzed by a Shimadzu LC Nexera X2 UHPLC coupled with a QTRAP 5500 LC-MS/MS (AB Sciex). An ACQUITY UPLC BEH Amide analytic column (2.1×50 mm, 1.7 μ Corp., Milford, MA) was used for chromatographic separation. The mobile phase was (A) water with 10 mM ammonium acetate (pH 8.9) and (B) acetonitrile/water (95/5) with 10 mM ammonium acetate (pH 8.2) (All solvents were LC–MS Optima grade from Fisher Scientific). The total run time was 11 min. with a flow rate of 0.5 ml/min. and an injection volume of 5 μ The gradient elution was 95–61% B in 6 min, 61–44% B at 8 min, 61–27% B at 8.2 min, and 27–95% B at 9 min. The column was equilibrated with 95% B at the end of each run. The source and collision gas was N_2_. The ion source conditions in positive and negative mode were: curtain gas (CUR) = 25 psi, collision gas (CAD) = high, ion spray voltage (IS) = 3800/-3800 volts, temperature (TEM) = 500 °C, ion source gas 1 (GS1) = 50 psi, and ion source gas 2 (GS2) = 40 psi. Each metabolite was tuned with standards for optimal transitions. D4-nicotinamide (Cambridge Isotope Laboratories, Tewksbury, MA) was used as an internal standard. The extracted MRM peaks were integrated using MultiQuant 3.0.3 software (AB Sciex, Concord, ON, CA).

### Statistical Analyses

Sample number (n) is defined as number of animals per genotype. Specific statistical tests and sample sizes are indicated in figure legends. Where applicable, p-value adjustments for multiple comparisons were performed and indicated, and reported as ‘q’ values. Across all figures, statistical significance is defined as p<0.05 (or q<0.05, where applicable). Experimenters were not blinded to treatments.

## ACKNOWLEDGEMENTS

We thank Dr. Laura Conforti for the *Nmnat1*^fl/fl^ animals, and current and past members of the Kolandaivelu lab for constant support. This work was supported by bridge funding to SK and the National Institutes of Health (RO1EY028959 to SK).

## REFERENCES

Agathocleous, Michalis, and William A. Harris. “Metabolism in Physiological Cell Proliferation and Differentiation.” Trends in Cell Biology 23, no. 10 (October 2013): 484–92. https://doi.org/10.1016/j.tcb.2013.05.004.

Aldiri, Issam, Beisi Xu, Lu Wang, Xiang Chen, Daniel Hiler, Lyra Griffiths, Marc Valentine, et al. “The Dynamic Epigenetic Landscape of the Retina During Development, Reprogramming, and Tumorigenesis.” Neuron 94, no. 3 (May 2017): 550–568.e10. https://doi.org/10.1016/j.neuron.2017.04.022.

Aleman, Tomas S., Katherine E. Uyhazi, Leona W. Serrano, Vidyullatha Vasireddy, Scott J. Bowman, Michael J. Ammar, Denise J. Pearson, Albert M. Maguire, and Jean Bennett. “RDH12 Mutations Cause a Severe Retinal Degeneration With Relatively Spared Rod Function.” Investigative Opthalmology & Visual Science 59, no. 12 (October 1, 2018): 5225. https://doi.org/10.1167/iovs.18-24708.

Bedoni, N., Quinodoz, M., Pinelli, M., Cappuccio, G., Torella, A., Nigro, V., Testa, F., Simonelli, F., TUDP (Telethon Undiagnosed Disease Program), Corton, M., Lualdi, S., Lanza, F., Morana, G., Ayuso, C., Di Rocco, M., Filocamo, M., Banfi, S., Brunetti-Pierri, N., Superti-Furga, A., & Rivolta, C. (2020). An Alu-mediated duplication in NMNAT1, involved in NAD biosynthesis, causes a novel syndrome, SHILCA, affecting multiple tissues and organs. Human Molecular Genetics, 29(13), 2250–2260. https://doi.org/10.1093/hmg/ddaa112

Bennett, Lea D., Martin Klein, Finny T. John, Bojana Radojevic, Kaylie Jones, and David G. Birch. “Disease Progression in Patients with Autosomal Dominant Retinitis Pigmentosa Due to a Mutation in Inosine Monophosphate Dehydrogenase 1 (IMPDH1).” Translational Vision Science & Technology 9, no. 5 (April 23, 2020): 14. https://doi.org/10.1167/tvst.9.5.14.

Bowne, Sara J., Lori S. Sullivan, Sarah E. Mortimer, Lizbeth Hedstrom, Jingya Zhu, Catherine J. Spellicy, Anisa I. Gire, et al. “Spectrum and Frequency of Mutations in IMPDH1 Associated with Autosomal Dominant Retinitis Pigmentosa and Leber Congenital Amaurosis.” Investigative Opthalmology & Visual Science 47, no. 1 (January 1, 2006): 34. https://doi.org/10.1167/iovs.05-0868.

Brazill, Jennifer M, Chong Li, Yi Zhu, and R Grace Zhai. “NMNAT: It’s an NAD + Synthase… It’s a Chaperone… It’s a Neuroprotector.” Current Opinion in Genetics & Development 44 (June 2017): 156–62. https://doi.org/10.1016/j.gde.2017.03.014.

Brzezinski, Joseph A, and Thomas A Reh. “Photoreceptor Cell Fate Specification in Vertebrates,” 2015, 11.

Cambronne, X. A., M. L. Stewart, D. Kim, A. M. Jones-Brunette, R. K. Morgan, D. L. Farrens, M. S. Cohen, and R. H. Goodman. “Biosensor Reveals Multiple Sources for Mitochondrial NAD+.” Science 352, no. 6292 (June 17, 2016): 1474–77. https://doi.org/10.1126/science.aad5168.

Cambronne, Xiaolu A., and W. Lee Kraus. “Location, Location, Location: Compartmentalization of NAD+ Synthesis and Functions in Mammalian Cells.” Trends in Biochemical Sciences 45, no. 10 (October 2020): 858–73. https://doi.org/10.1016/j.tibs.2020.05.010.

Cantó, Carles, Keir J. Menzies, and Johan Auwerx. “NAD+ Metabolism and the Control of Energy Homeostasis: A Balancing Act between Mitochondria and the Nucleus.” Cell Metabolism 22, no. 1 (July 2015): 31–53. https://doi.org/10.1016/j.cmet.2015.05.023.

Carty, Michael, Jay Kearney, Katharine A. Shanahan, Emily Hams, Ryoichi Sugisawa, Dympna Connolly, Ciara G. Doran, et al. “Cell Survival and Cytokine Release after Inflammasome Activation Is Regulated by the Toll-IL-1R Protein SARM.” Immunity 50, no. 6 (June 2019): 1412–1424.e6. https://doi.org/10.1016/j.immuni.2019.04.005.

Chang, J., B. Zhang, H. Heath, N. Galjart, X. Wang, and J. Milbrandt. “Nicotinamide Adenine Dinucleotide (NAD)-Regulated DNA Methylation Alters CCCTC-Binding Factor (CTCF)/Cohesin Binding and Transcription at the BDNF Locus.” Proceedings of the National Academy of Sciences 107, no. 50 (December 14, 2010): 21836–41. https://doi.org/10.1073/pnas.1002130107.

Chiang, Pei-Wen, Juan Wang, Yang Chen, Quan Fu, Jing Zhong, Yanhua Chen, Xin Yi, et al. “Exome Sequencing Identifies NMNAT1 Mutations as a Cause of Leber Congenital Amaurosis.” Nature Genetics 44, no. 9 (September 2012): 972–74. https://doi.org/10.1038/ng.2370.

Chinchore, Yashodhan, Tedi Begaj, David Wu, Eugene Drokhlyansky, and Constance L Cepko. “Glycolytic Reliance Promotes Anabolism in Photoreceptors.” ELife 6 (June 9, 2017): e25946. https://doi.org/10.7554/eLife.25946.

Christiansen, J. R., S. Kolandaivelu, M. O. Bergo, and V. Ramamurthy. “RAS-Converting Enzyme 1-Mediated Endoproteolysis Is Required for Trafficking of Rod Phosphodiesterase 6 to Photoreceptor Outer Segments.” Proceedings of the National Academy of Sciences 108, no. 21 (May 24, 2011): 8862–66. https://doi.org/10.1073/pnas.1103627108.

Conforti, Laura, Lucie Janeckova, Diana Wagner, Francesca Mazzola, Lucia Cialabrini, Michele Di Stefano, Giuseppe Orsomando, et al. “Reducing Expression of NAD+ Synthesizing Enzyme NMNAT1 Does Not Affect the Rate of Wallerian Degeneration.” The FEBS Journal 278, no. 15 (2011): 2666–79. https://doi.org/10.1111/j.1742-4658.2011.08193.x.

Coppieters, Frauke, Anne Laure Todeschini, Takuro Fujimaki, Annelot Baert, Marieke Bruyne, Caroline Cauwenbergh, Hannah Verdin, et al. “Hidden Genetic Variation in LCA9- Associated Congenital Blindness Explained by 5′UTR Mutations and Copy-Number Variations of NMNAT1.” Human Mutation 36, no. 12 (December 2015): 1188–96. https://doi.org/10.1002/humu.22899.

Daum, Janine M, Özkan Keles, Sjoerd JB Holwerda, Hubertus Kohler, Filippo M Rijli, Michael Stadler, and Botond Roska. “The Formation of the Light-Sensing Compartment of Cone Photoreceptors Coincides with a Transcriptional Switch.” ELife 6 (November 6, 2017): e31437. https://doi.org/10.7554/eLife.31437.

Donovan, M, and T G Cotter. “Caspase-Independent Photoreceptor Apoptosis in Vivo and Differential Expression of Apoptotic Protease Activating Factor-1 and Caspase-3 during Retinal Development.” Cell Death & Differentiation 9, no. 11 (November 2002): 1220–31. https://doi.org/10.1038/sj.cdd.4401105.

Donovan, Maryanne, Francesca Doonan, and Thomas G. Cotter. “Decreased Expression of Pro-Apoptotic Bcl-2 Family Members during Retinal Development and Differential Sensitivity to Cell Death.” Developmental Biology 291, no. 1 (March 2006): 154–69. https://doi.org/10.1016/j.ydbio.2005.12.026.

Doonan, Francesca, Maryanne Donovan, and Thomas G. Cotter. “Caspase-Independent Photoreceptor Apoptosis in Mouse Models of Retinal Degeneration.” The Journal of Neuroscience 23, no. 13 (July 2, 2003): 5723–31. https://doi.org/10.1523/JNEUROSCI.23-13-05723.2003.

Du, Jianhai, Haiwei Gu, Sally J Turner, Danijel Djukovic, Daniel Raftery, and James B Hurley. “Metabolite Profiles of Rod Photoreceptor Cell Death in Mouse Retinal Degeneration Models.” Investigative Ophthalmology & Visual Science 55, no. 13 (April 30, 2014): 4374–4374.

Du, Jianhai, Jonathan D. Linton, and James B. Hurley. “Probing Metabolism in the Intact Retina Using Stable Isotope Tracers.” In Methods in Enzymology, 561:149–70. Elsevier, 2015. https://doi.org/10.1016/bs.mie.2015.04.002.

Dvoriantchikova, Galina, Seemungal, Rajeev, and Dmitry Ivanov. “DNA Methylation Dynamics During the Differentiation of Retinal Progenitor Cells Into Retinal Neurons Reveal a Role for the DNA Demethylation Pathway.” Frontiers in Molecular Neuroscience 12 (2019): 9.

Eblimit, Aiden, Smriti Agrawal Zaneveld, Wei Liu, Kandace Thomas, Keqing Wang, Yumei Li, Graeme Mardon, and Rui Chen. “NMNAT1 E257K Variant, Associated with Leber Congenital Amaurosis (LCA9), Causes a Mild Retinal Degeneration Phenotype.” Experimental Eye Research 173 (August 2018): 32–43. https://doi.org/10.1016/j.exer.2018.04.010.

Etingof, R N. “De Novo Biosynthesis of Purines in the Retina: Evolutionary Aspects” 37, no. 1 (2001): 10.

Falk, Marni J, Qi Zhang, Eiko Nakamaru-Ogiso, Chitra Kannabiran, Zoe Fonseca-Kelly, Christina Chakarova, Isabelle Audo, et al. “NMNAT1 Mutations Cause Leber Congenital Amaurosis.” Nature Genetics 44, no. 9 (September 2012): 1040–45. https://doi.org/10.1038/ng.2361.

Figley, Matthew D., Weixi Gu, Jeffrey D. Nanson, Yun Shi, Yo Sasaki, Katie Cunnea, Alpeshkumar K. Malde, et al. “SARM1 Is a Metabolic Sensor Activated by an Increased NMN/NAD+ Ratio to Trigger Axon Degeneration.” Neuron, March 2021, S0896627321000830. https://doi.org/10.1016/j.neuron.2021.02.009.

Finding of Rare Disease Genes (FORGE) Canada Consortium, Robert K Koenekoop, Hui Wang, Jacek Majewski, Xia Wang, Irma Lopez, Huanan Ren, et al. “Mutations in NMNAT1 Cause Leber Congenital Amaurosis and Identify a New Disease Pathway for Retinal Degeneration.” Nature Genetics 44, no. 9 (September 2012): 1035–39. https://doi.org/10.1038/ng.2356.

Furuta, Yasuhide, Oleg Lagutin, Brigid L M Hogan, and Guillermo C Oliver. “Retina- and Ventral Forebrain-specific Cre Recombinase Activity in Transgenic Mice,” n.d., 3.

Greenwald, Scott H, Emily E Brown, Michael J Scandura, Erin Hennessey, Raymond Farmer, Jianhai Du, Yekai Wang, and Eric A Pierce. “Mutant NMNAT1 Leads to a Retina-Specific Decrease of NAD+ Accompanied by Increased Poly(ADP-Ribose) in a Mouse Model of NMNAT1-Associated Retinal Degeneration.” Human Molecular Genetics, March 11, 2021, ddab070. https://doi.org/10.1093/hmg/ddab070.

Greenwald, Scott H., Emily E. Brown, Michael J. Scandura, Erin Hennessey, Raymond Farmer, Basil S. Pawlyk, Ru Xiao, Luk H. Vandenberghe, and Eric A. Pierce. “Gene Therapy Preserves Retinal Structure and Function in a Mouse Model of NMNAT1-Associated Retinal Degeneration.” Molecular Therapy - Methods & Clinical Development 18 (September 2020): 582–94. https://doi.org/10.1016/j.omtm.2020.07.003.

Greenwald, Scott H., Jeremy R. Charette, Magdalena Staniszewska, Lan Ying Shi, Steve D.M. Brown, Lisa Stone, Qin Liu, et al. “Mouse Models of NMNAT1-Leber Congenital Amaurosis (LCA9) Recapitulate Key Features of the Human Disease.” The American Journal of Pathology 186, no. 7 (July 2016): 1925–38. https://doi.org/10.1016/j.ajpath.2016.03.013.

Grenell, A., Wang, Y., Yam, M., Swarup, A., Dilan, T. L., Hauer, A., Linton, J. D., Philp, N. J., Gregor, E., Zhu, S., Shi, Q., Murphy, J., Guan, T., Lohner, D., Kolandaivelu, S., Ramamurthy, V., Goldberg, A. F. X., Hurley, J. B., & Du, J. (2019). Loss of MPC1 reprograms retinal metabolism to impair visual function. Proceedings of the National Academy of Sciences, 116(9), 3530–3535. https://doi.org/10.1073/pnas.1812941116

Gutierrez, Kimberley D., Michael A. Davis, Brian P. Daniels, Tayla M. Olsen, Pooja Ralli-Jain, Stephen W. G. Tait, Michael Gale, and Andrew Oberst. “MLKL Activation Triggers NLRP3-Mediated Processing and Release of IL-1β Independently of Gasdermin-D.” The Journal of Immunology 198, no. 5 (March 1, 2017): 2156–64. https://doi.org/10.4049/jimmunol.1601757.

Houtkooper, Riekelt H., Carles Cantó, Ronald J. Wanders, and Johan Auwerx. “The Secret Life of NAD +□: An Old Metabolite Controlling New Metabolic Signaling Pathways.” Endocrine Reviews 31, no. 2 (April 2010): 194–223. https://doi.org/10.1210/er.2009-0026.

Kanow, Mark A, Michelle M Giarmarco, Connor SR Jankowski, Kristine Tsantilas, Abbi L Engel, Jianhai Du, Jonathan D Linton, et al. “Biochemical Adaptations of the Retina and Retinal Pigment Epithelium Support a Metabolic Ecosystem in the Vertebrate Eye.” ELife 6 (September 13, 2017): e28899. https://doi.org/10.7554/eLife.28899.

Kayagaki, Nobuhiko, Bettina L. Lee, Irma B. Stowe, Opher S. Kornfeld, Karen O’Rourke, Kathleen M. Mirrashidi, Benjamin Haley, et al. “IRF2 Transcriptionally Induces GSDMD Expression for Pyroptosis.” Science Signaling 12, no. 582 (May 21, 2019): eaax4917. https://doi.org/10.1126/scisignal.aax4917.

Kim, Daehwan, Ben Langmead, and Steven L Salzberg. “HISAT: A Fast Spliced Aligner with Low Memory Requirements.” Nature Methods 12, no. 4 (April 2015): 357–60. https://doi.org/10.1038/nmeth.3317.

Kumaran, Neruban, Anthony T Moore, Richard G Weleber, and Michel Michaelides. “Leber Congenital Amaurosis/Early-Onset Severe Retinal Dystrophy: Clinical Features, Molecular Genetics and Therapeutic Interventions.” British Journal of Ophthalmology 101, no. 9 (September 2017): 1147–54. https://doi.org/10.1136/bjophthalmol-2016-309975.

Kuribayashi, Hiroshi, Yukihiro Baba, Toshiro Iwagawa, Eisuke Arai, Akira Murakami, and Sumiko Watanabe. “Roles of Nmnat1 in the Survival of Retinal Progenitors through the Regulation of Pro-Apoptotic Gene Expression via Histone Acetylation.” Cell Death & Disease 9, no. 9 (August 30, 2018). https://doi.org/10.1038/s41419-018-0907-0.

Lin, Jonathan B., Shunsuke Kubota, Norimitsu Ban, Mitsukuni Yoshida, Andrea Santeford, Abdoulaye Sene, Rei Nakamura, et al. “NAMPT-Mediated NAD + Biosynthesis Is Essential for Vision In Mice.” Cell Reports 17, no. 1 (September 2016): 69–85. https://doi.org/10.1016/j.celrep.2016.08.073.

Love, Michael I, Wolfgang Huber, and Simon Anders. “Moderated Estimation of Fold Change and Dispersion for RNA-Seq Data with DESeq2.” Genome Biology 15, no. 12 (December 2014): 550. https://doi.org/10.1186/s13059-014-0550-8.

Lundt, Samuel, Nannan Zhang, Jun-Liszt Li, Zhe Zhang, Li Zhang, Xiaowan Wang, Ruisi Bao, et al. “Metabolomic and Transcriptional Profiling Reveals Bioenergetic Stress and Activation of Cell Death and Inflammatory Pathways in Vivo after Neuronal Deletion of NAMPT.” Journal of Cerebral Blood Flow & Metabolism, February 9, 2021, 0271678X2199262. https://doi.org/10.1177/0271678X21992625.

McKenzie, Brienne A. “Fiery Cell Death: Pyroptosis in the Central Nervous System.” Trends in Neurosciences, n.d., 19.

Miao, Edward A., Jayant V. Rajan, and Alan Aderem. “Caspase-1-Induced Pyroptotic Cell Death: Caspase-1-Induced Pyroptotic Cell Death.” Immunological Reviews 243, no. 1 (September 2011): 206–14. https://doi.org/10.1111/j.1600-065X.2011.01044.x.

Mori, Valerio, Adolfo Amici, Francesca Mazzola, Michele Di Stefano, Laura Conforti, Giulio Magni, Silverio Ruggieri, Nadia Raffaelli, and Giuseppe Orsomando. “Metabolic Profiling of Alternative NAD Biosynthetic Routes in Mouse Tissues.” Edited by Valerie de Crécy-Lagard. PLoS ONE 9, no. 11 (November 25, 2014): e113939. https://doi.org/10.1371/journal.pone.0113939.

Mukherjee, Piyali, Clayton W. Winkler, Katherine G. Taylor, Tyson A. Woods, Vinod Nair, Burhan A. Khan, and Karin E. Peterson. “SARM1, Not MyD88, Mediates TLR7/TLR9-Induced Apoptosis in Neurons.” The Journal of Immunology 195, no. 10 (November 15, 2015): 4913–21. https://doi.org/10.4049/jimmunol.1500953.

Müller, Tammo, Christin Dewitz, Jessica Schmitz, Anna Sophia Schröder, Jan Hinrich Bräsen, Brent R. Stockwell, James M. Murphy, Ulrich Kunzendorf, and Stefan Krautwald. “Necroptosis and Ferroptosis Are Alternative Cell Death Pathways That Operate in Acute Kidney Failure.” Cellular and Molecular Life Sciences 74, no. 19 (October 2017): 3631–45. https://doi.org/10.1007/s00018-017-2547-4.

Murakami, Y., H. Matsumoto, M. Roh, J. Suzuki, T. Hisatomi, Y. Ikeda, J. W. Miller, and D. G. Vavvas. “Receptor Interacting Protein Kinase Mediates Necrotic Cone but Not Rod Cell Death in a Mouse Model of Inherited Degeneration.” Proceedings of the National Academy of Sciences 109, no. 36 (September 4, 2012): 14598–603. https://doi.org/10.1073/pnas.1206937109.

Nash, Benjamin M., Richard Symes, Himanshu Goel, Marcel E. Dinger, Bruce Bennetts, John R. Grigg, and Robyn V. Jamieson. “NMNAT1 Variants Cause Cone and Cone-Rod Dystrophy.” European Journal of Human Genetics 26, no. 3 (March 2018): 428–33. https://doi.org/10.1038/s41431-017-0029-7.

Navas, Lola E., and Amancio Carnero. “NAD+ Metabolism, Stemness, the Immune Response, and Cancer.” Signal Transduction and Targeted Therapy 6, no. 1 (December 2021): 2. https://doi.org/10.1038/s41392-020-00354-w.

Newton, K., D. L. Dugger, K. E. Wickliffe, N. Kapoor, M. C. de Almagro, D. Vucic, L. Komuves, et al. “Activity of Protein Kinase RIPK3 Determines Whether Cells Die by Necroptosis or Apoptosis.” Science 343, no. 6177 (March 21, 2014): 1357–60. https://doi.org/10.1126/science.1249361.

Ng, Soo Khai, John PM Wood, Glyn Chidlow, Guoge Han, Thaksaon Kittipassorn, Daniel J Peet, and Robert J Casson. “Cancer-like Metabolism of the Mammalian Retina: Mammalian Retina Metabolism.” Clinical & Experimental Ophthalmology 43, no. 4 (May 2015): 367–76. https://doi.org/10.1111/ceo.12462.

Nikiforov, Andrey, Veronika Kulikova, and Mathias Ziegler. “The Human NAD Metabolome: Functions, Metabolism and Compartmentalization.” Critical Reviews in Biochemistry and Molecular Biology 50, no. 4 (July 4, 2015): 284–97. https://doi.org/10.3109/10409238.2015.1028612.

Oakey, Lucy A., Rachel S. Fletcher, Yasir S. Elhassan, David M. Cartwright, Craig L. Doig, Antje Garten, Alpesh Thakker, et al. “Metabolic Tracing Reveals Novel Adaptations to Skeletal Muscle Cell Energy Production Pathways in Response to NAD + Depletion.” Wellcome Open Research 3 (2018): 147. https://doi.org/10.12688/wellcomeopenres.14898.2.

Perrault, Isabelle, Sylvain Hanein, Xavier Zanlonghi, Valérie Serre, Michael Nicouleau, Sabine Defoort-Delhemmes, Nathalie Delphin, et al. “Mutations in NMNAT1 Cause Leber Congenital Amaurosis with Early-Onset Severe Macular and Optic Atrophy.” Nature Genetics 44, no. 9 (September 2012): 975–77. https://doi.org/10.1038/ng.2357.

Pertea, Mihaela, Daehwan Kim, Geo M Pertea, Jeffrey T Leek, and Steven L Salzberg. “Transcript-Level Expression Analysis of RNA-Seq Experiments with HISAT, StringTie and Ballgown.” Nature Protocols 11, no. 9 (August 11, 2016): 1650–67. https://doi.org/10.1038/nprot.2016.095.

Pertea, Mihaela, Geo M Pertea, Corina M Antonescu, Tsung-Cheng Chang, Joshua T Mendell, and Steven L Salzberg. “StringTie Enables Improved Reconstruction of a Transcriptome from RNA-Seq Reads.” Nature Biotechnology 33, no. 3 (March 2015): 290–95. https://doi.org/10.1038/nbt.3122.

Preyat, Nicolas, and Oberdan Leo. “Complex Role of Nicotinamide Adenine Dinucleotide in the Regulation of Programmed Cell Death Pathways.” Biochemical Pharmacology 101 (February 2016): 13–26. https://doi.org/10.1016/j.bcp.2015.08.110.

Raeisossadati, Reza, Merari F. R. Ferrari, Alexandre Hiroaki Kihara, Issam AlDiri, and Jeffrey M. Gross. “Epigenetic Regulation of Retinal Development.” Epigenetics & Chromatin 14, no. 1 (December 2021): 11. https://doi.org/10.1186/s13072-021-00384-w.

Rhee, K-D, J Yu, C Y Zhao, G Fan, and X-J Yang. “Dnmt1-Dependent DNA Methylation Is Essential for Photoreceptor Terminal Differentiation and Retinal Neuron Survival.” Cell Death & Disease 3, no. 11 (November 2012): e427–e427. https://doi.org/10.1038/cddis.2012.165.

Sanman, Laura E, Yu Qian, Nicholas A Eisele, Tessie M Ng, Wouter A van der Linden, Denise M Monack, Eranthie Weerapana, and Matthew Bogyo. “Disruption of Glycolytic Flux Is a Signal for Inflammasome Signaling and Pyroptotic Cell Death.” ELife 5 (March 24, 2016): e13663. https://doi.org/10.7554/eLife.13663.

Sasaki, Yo, Thomas M. Engber, Robert O. Hughes, Matthew D. Figley, Tong Wu, Todd Bosanac, Rajesh Devraj, Jeffrey Milbrandt, Raul Krauss, and Aaron DiAntonio. “CADPR Is a Gene Dosage-Sensitive Biomarker of SARM1 Activity in Healthy, Compromised, and Degenerating Axons.” Experimental Neurology 329 (July 2020): 113252. https://doi.org/10.1016/j.expneurol.2020.113252.

Sasaki, Yo, Hiroki Kakita, Shunsuke Kubota, Abdoulaye Sene, Tae Jun Lee, Norimitsu Ban, Zhenyu Dong, et al. “SARM1 Depletion Rescues NMNAT1-Dependent Photoreceptor Cell Death and Retinal Degeneration.” ELife 9 (October 27, 2020): e62027. https://doi.org/10.7554/eLife.62027.

Sasaki, Yo, Zachary Margolin, Benjamin Borgo, James J. Havranek, and Jeffrey Milbrandt. “Characterization of Leber Congenital Amaurosis-Associated NMNAT1 Mutants.” Journal of Biological Chemistry 290, no. 28 (July 2015): 17228–38. https://doi.org/10.1074/jbc.M115.637850.

Sato, K., S. Li, W. C. Gordon, J. He, G. I. Liou, J. M. Hill, G. H. Travis, N. G. Bazan, and M. Jin. “Receptor Interacting Protein Kinase-Mediated Necrosis Contributes to Cone and Rod Photoreceptor Degeneration in the Retina Lacking Interphotoreceptor Retinoid-Binding Protein.” Journal of Neuroscience 33, no. 44 (October 30, 2013): 17458–68. https://doi.org/10.1523/JNEUROSCI.1380-13.2013.

Seritrakul, Pawat, and Jeffrey M. Gross. “Tet-Mediated DNA Hydroxymethylation Regulates Retinal Neurogenesis by Modulating Cell-Extrinsic Signaling Pathways.” Edited by Wolf Reik. PLOS Genetics 13, no. 9 (September 19, 2017): e1006987. https://doi.org/10.1371/journal.pgen.1006987.

Singh, Ratnesh K., Ramya K. Mallela, Abigail Hayes, Nicholas R. Dunham, Morgan E. Hedden, Raymond A. Enke, Robert N. Fariss, Hal Sternberg, Michael D. West, and Igor O. Nasonkin. “Dnmt1, Dnmt3a and Dnmt3b Cooperate in Photoreceptor and Outer Plexiform Layer Development in the Mammalian Retina.” Experimental Eye Research 159 (June 2017): 132–46. https://doi.org/10.1016/j.exer.2016.11.014.

Sinha, Tirthankar, Jianhai Du, Mustafa S. Makia, James B. Hurley, Muna I. Naash, and Muayyad R. Al-Ubaidi. “Absence of Retbindin Blocks Glycolytic Flux, Disrupts Metabolic Homeostasis, and Leads to Photoreceptor Degeneration.” Proceedings of the National Academy of Sciences 118, no. 6 (February 9, 2021): e2018956118. https://doi.org/10.1073/pnas.2018956118.

Song, Tanjing, Leixiang Yang, Neha Kabra, Lihong Chen, John Koomen, Eric B. Haura, and Jiandong Chen. “The NAD+ Synthesis Enzyme Nicotinamide Mononucleotide Adenylyltransferase (NMNAT1) Regulates Ribosomal RNA Transcription.” Journal of Biological Chemistry 288, no. 29 (July 19, 2013): 20908–17. https://doi.org/10.1074/jbc.M113.470302.

Svoboda, Petr, Edita Krizova, Sarka Sestakova, Kamila Vapenkova, Zdenek Knejzlik, Silvie Rimpelova, Diana Rayova, et al. “Nuclear Transport of Nicotinamide Phosphoribosyltransferase Is Cell Cycle–Dependent in Mammalian Cells, and Its Inhibition Slows Cell Growth.” Journal of Biological Chemistry 294, no. 22 (May 2019): 8676–89. https://doi.org/10.1074/jbc.RA118.003505.

Swanson, Karen V., Meng Deng, and Jenny P.-Y. Ting. “The NLRP3 Inflammasome: Molecular Activation and Regulation to Therapeutics.” Nature Reviews Immunology 19, no. 8 (August 2019): 477–89. https://doi.org/10.1038/s41577-019-0165-0.

Swaroop, Anand, Douglas Kim, and Douglas Forrest. “Transcriptional Regulation of Photoreceptor Development and Homeostasis in the Mammalian Retina.” Nature Reviews Neuroscience 11, no. 8 (August 2010): 563–76. https://doi.org/10.1038/nrn2880.

Vandenabeele, Peter, Lorenzo Galluzzi, Tom Vanden Berghe, and Guido Kroemer. “Molecular Mechanisms of Necroptosis: An Ordered Cellular Explosion.” Nature Reviews Molecular Cell Biology 11, no. 10 (October 2010): 700–714. https://doi.org/10.1038/nrm2970.

Wang, Xiaolin, Yu Fang, Rongsheng Liao, and Tao Wang. “Targeted Deletion of Nmnat1 in Mouse Retina Leads to Early Severe Retinal Dystrophy.” Preprint. Developmental Biology, October 29, 2017. https://doi.org/10.1101/210757.

Yam, M., Engel, A. L., Wang, Y., Zhu, S., Hauer, A., Zhang, R., Lohner, D., Huang, J., Dinterman, M., Zhao, C., Chao, J. R., & Du, J. (2019). Proline mediates metabolic communication between retinal pigment epithelial cells and the retina. Journal of Biological Chemistry, 294(26), 10278–10289. https://doi.org/10.1074/jbc.RA119.007983

Zhang, Rui, Weiyong Shen, Jianhai Du, and Mark C. Gillies. “Selective Knockdown of Hexokinase 2 in Rods Leads to Age-Related Photoreceptor Degeneration and Retinal Metabolic Remodeling.” Cell Death & Disease 11, no. 10 (October 2020): 885. https://doi.org/10.1038/s41419-020-03103-7.

Zhang, Tong, Jhoanna G. Berrocal, Jie Yao, Michelle E. DuMond, Raga Krishnakumar, Donald D. Ruhl, Keun Woo Ryu, Matthew J. Gamble, and W. Lee Kraus. “Regulation of Poly(ADP-Ribose) Polymerase-1-Dependent Gene Expression through Promoter-Directed Recruitment of a Nuclear NAD+ Synthase*.” Journal of Biological Chemistry 287, no. 15 (April 2012): 12405–16. https://doi.org/10.1074/jbc.M111.304469.

Zhao, Zhi Ying, Xu Jie Xie, Wan Hua Li, Jun Liu, Zhe Chen, Ben Zhang, Ting Li, et al. “A Cell-Permeant Mimetic of NMN Activates SARM1 to Produce Cyclic ADP-Ribose and Induce Non-Apoptotic Cell Death.” IScience 15 (May 31, 2019): 452–66. https://doi.org/10.1016/j.isci.2019.05.001.

Zhu, Siyan, Michelle Yam, Yekai Wang, Jonathan D. Linton, Allison Grenell, James B. Hurley, and Jianhai Du. “Impact of Euthanasia, Dissection and Postmortem Delay on Metabolic Profile in Mouse Retina and RPE/Choroid.” Experimental Eye Research 174 (September 2018): 113–20. https://doi.org/10.1016/j.exer.2018.05.032.

