## Supplemental Information for "Nuclear NAD^+^-biosynthetic enzyme NMNAT1 facilitates survival of developing retinal neurons"

David Sokolov[1]

Emily Sechrest[1,2]

Yekai Wang[1,3]

Connor Nevin[1]

Jianhai Du[1,3]

Saravanan Kolandaivelu[1,3,*]

**Affiliations**

[1]Department of Ophthalmology and Visual Sciences, Eye Institute, One Medical Center Drive, West Virginia University, Morgantown, WV, 26506-9193, USA

[2]Department of Pharmaceutical Sciences, One Medical Center Drive, West Virginia University, Morgantown, WV, 26506-9193, USA

[3]Department of Biochemistry, One Medical Center Drive, West Virginia University, Morgantown, WV, 26506-9193, USA

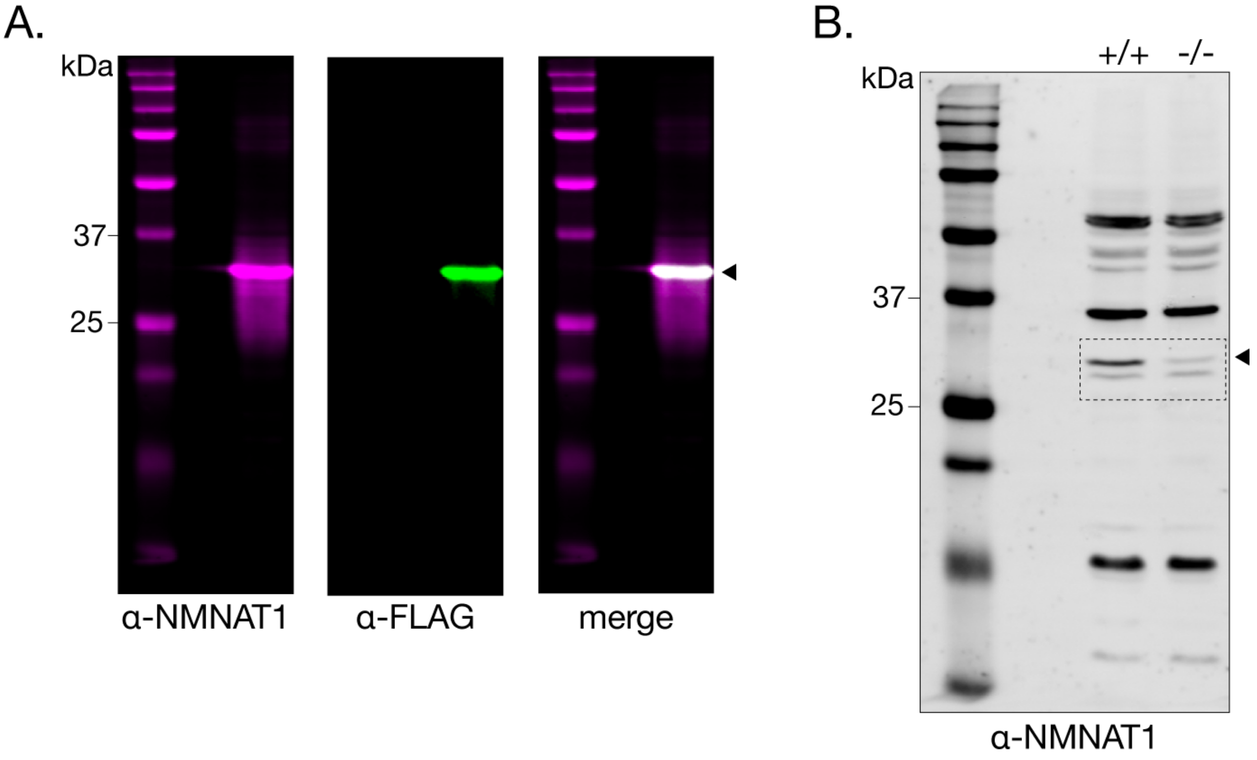


**Figure 1—Supplement 1. Validation of anti-NMNAT1 antibody in cell lines and retinal tissue.** (**A**) western blot of lysate of HEK293T cells transiently transfected with FLAG-tagged NMNAT1 demonstrating specificity of NMNAT1 antibody. (**B**) western blot from Figure 1C, with dotted line indicating crop area. Arrowheads denote bands corresponding to NMNAT1.

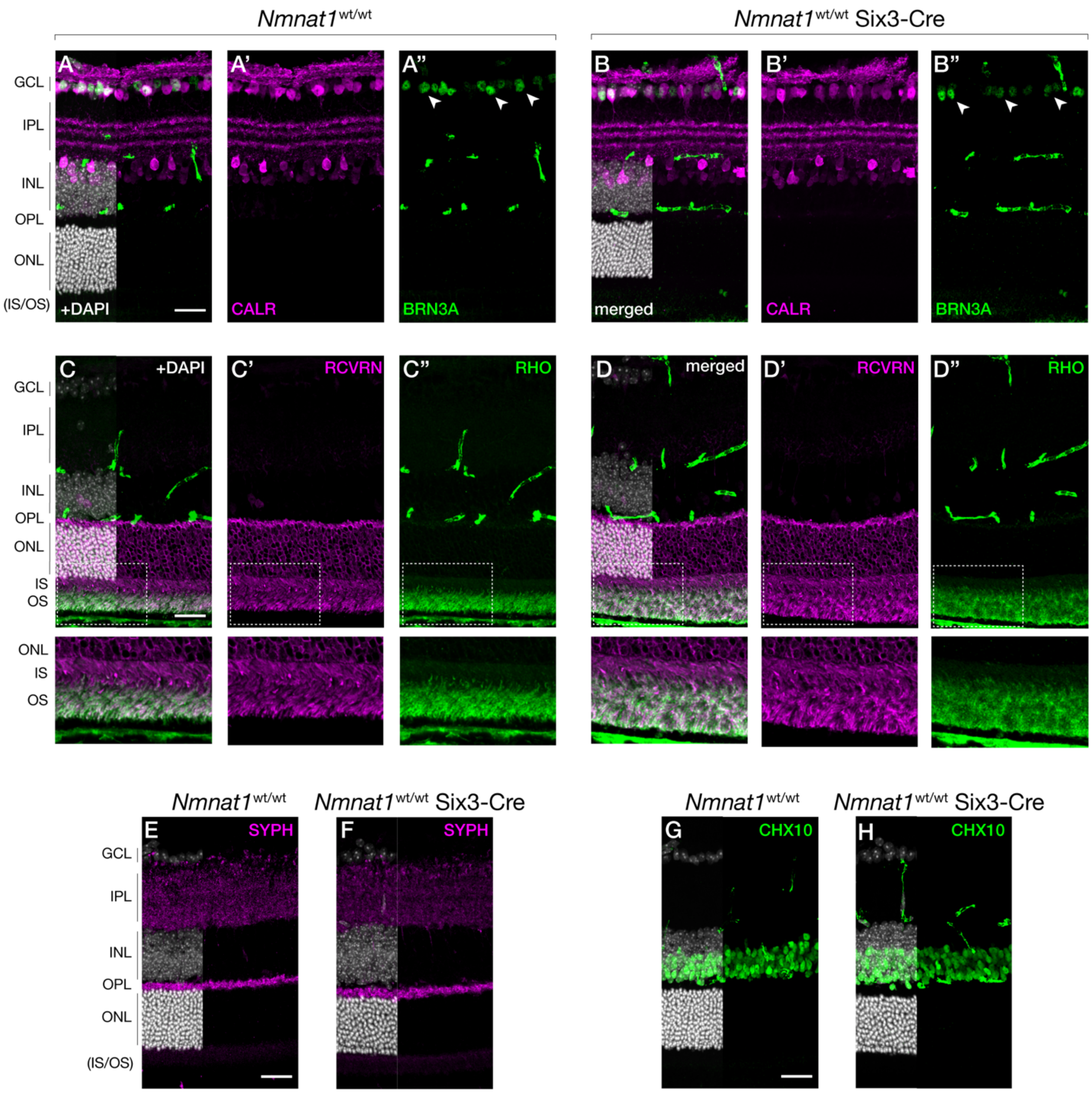


**Figure 2—Supplement 1. Six3-CRE does not cause obvious defects in the mature retina.** Representative retinal sections from P65 wild-type (*Nmnat1*$^{wt/wt}$) and Six3-Cre expressing (*Nmnat1*$^{wt/wt}$ *Six3-Cre*) mice labelled with antibodies against calretinin (CALR) and BRN3A (**A-B**), recoverin (RCVRN) and rhodopsin (RHO) (**C-D**), synaptophysin (SYPH) (**E,F**), or CHX10 (**G,H**). Corresponding zoom panels are indicated with dotted rectangles. n=3 biological replicates for all panels. Scale bars, 30 µm. Abbreviations: P, postnatal day; GCL, ganglion cell layer; IPL, inner plexiform layer; OPL, outer plexiform layer; INL, inner nuclear layer; ONL, outer nuclear layer; IS/OS, photoreceptor inner segment/outer segment layer.

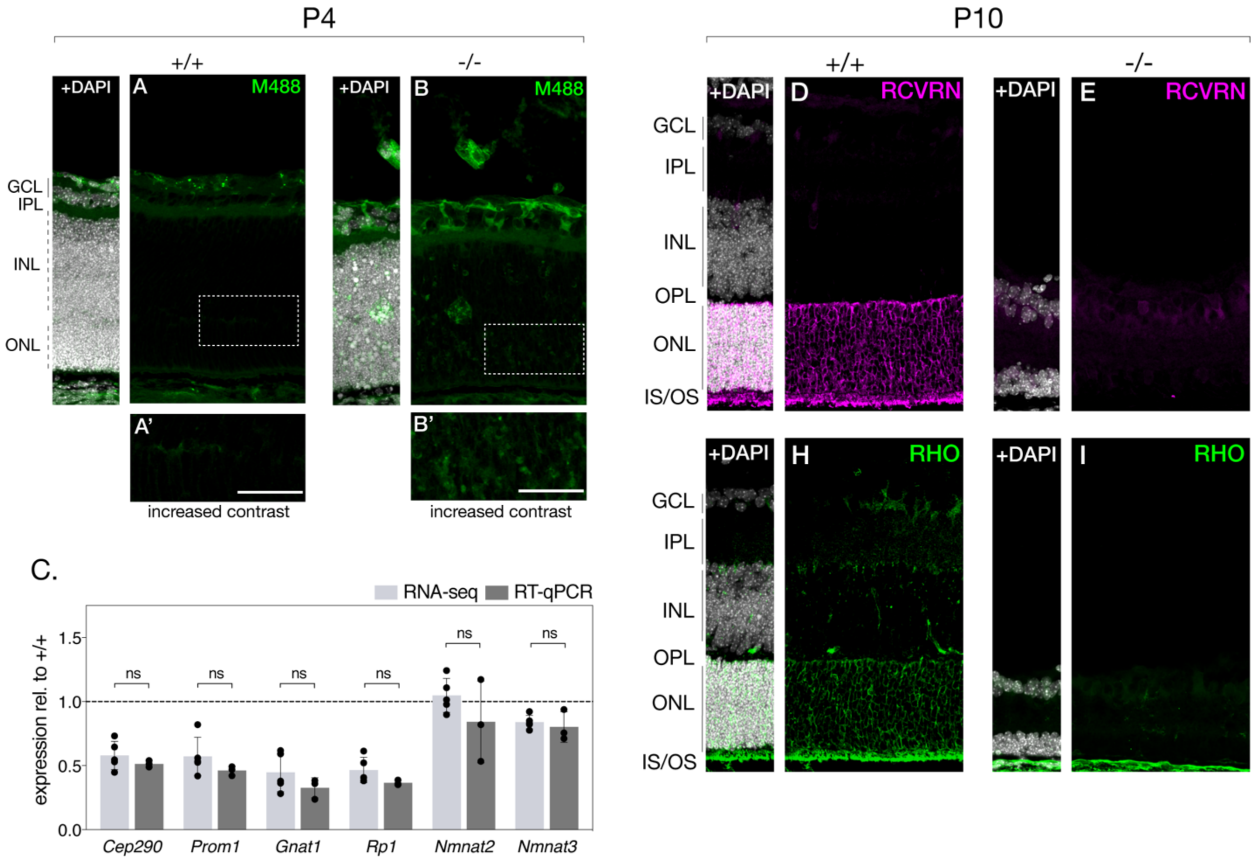


**Figure 3—Supplement 1.** **(A-B)** Representative retinal sections from P4 knockout (-/-) and control (+/+) mice labelled with anti-mouse 488 secondary antibody. (**C**) RT-qPCR validation of indicated genes in P4 RNA-sequencing dataset. Representative retinal sections from P10 knockout and control mice labelled with antibodies against recoverin (RCVRN) (**D**,**E**) and rhodopsin (RHO) (**H**,**I**) are also shown. Corresponding zoom panels are indicated with dotted rectangles. Scale bars, 30 µm. . Data is represented as mean ± SD. significance determined using Student's t-test.

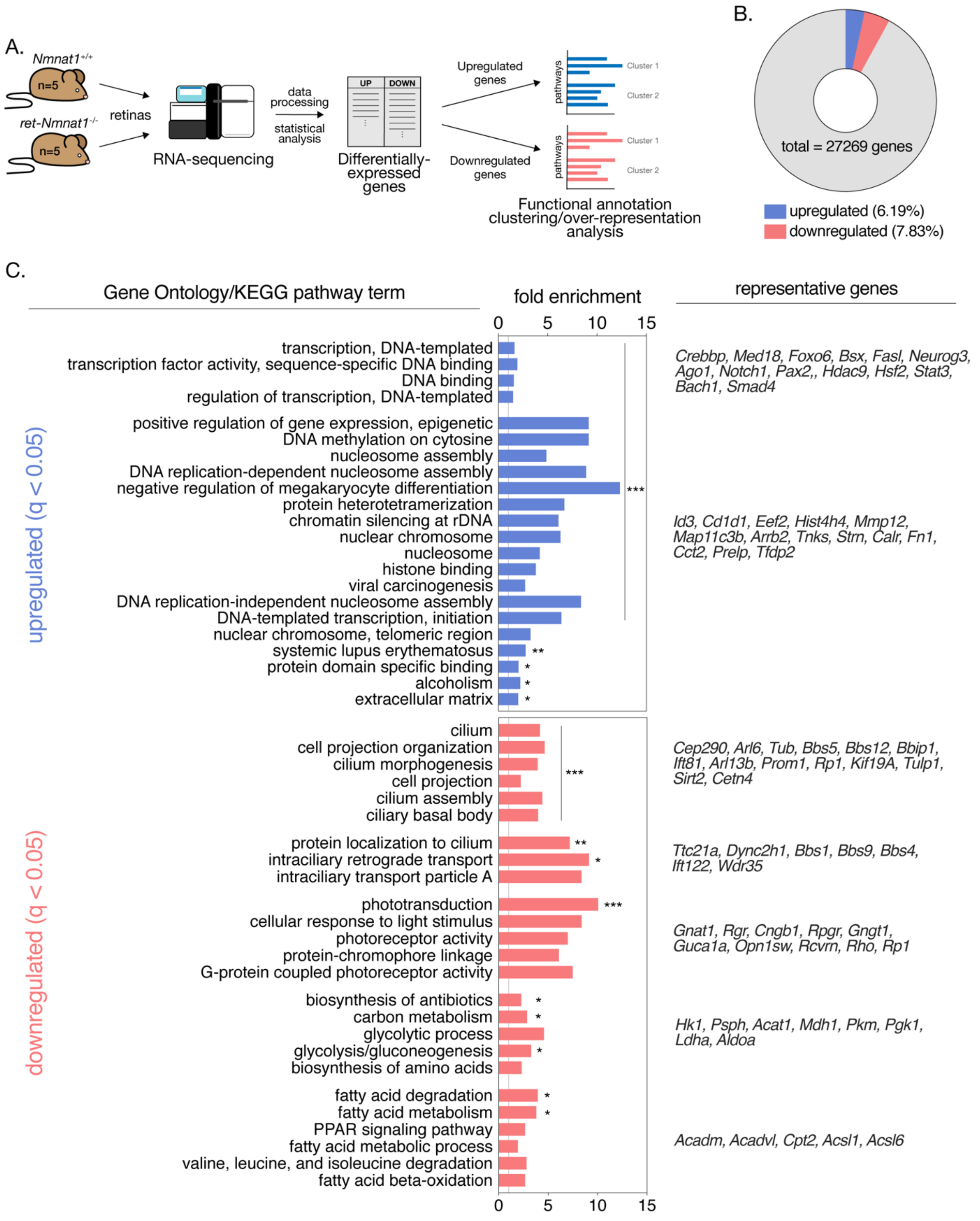
A.
Nmnat1+/+
n=5
ret-Nmnat1-/-
n=5
retinas
RNA-sequencing
data processing
statistical analysis
UP
DOWN
Differentially-expressed genes
Upregulated genes
Downregulated genes
pathways
Cluster 1
Cluster 2
Functional annotation clustering/over-representation analysis
B.
total = 27269 genes
upregulated (6.19%)
downregulated (7.83%)
C.
Gene Ontology/KEGG pathway term
fold enrichment
representative genes
0 5 10 15
upregulated (q < 0.05)
transcription, DNA-templated
transcription factor activity, sequence-specific DNA binding
DNA binding
regulation of transcription, DNA-templated
positive regulation of gene expression, epigenetic
DNA methylation on cytosine
nucleosome assembly
DNA replication-dependent nucleosome assembly
negative regulation of megakaryocyte differentiation
protein heterotetramerization
chromatin silencing at rDNA
nuclear chromosome
nucleosome
histone binding
viral carcinogenesis
DNA replication-independent nucleosome assembly
DNA-templated transcription, initiation
nuclear chromosome, telomeric region
systemic lupus erythematosus
protein domain specific binding
alcoholism
extracellular matrix
***
**
*
Crebbp, Med18, Foxo6, Bsx, Fasl, Neurog3, Ago1, Notch1, Pax2,, Hdac9, Hsf2, Stat3, Bach1, Smad4
Id3, Cd1d1, Eef2, Hist4h4, Mmp12, Map11c3b, Arrb2, Tnks, Strn, Calr, Fn1, Cct2, Prelp, Tfdp2
downregulated (q < 0.05)
cilium
cell projection organization
cilium morphogenesis
cell projection
cilium assembly
ciliary basal body
protein localization to cilium
intraciliary retrograde transport
intraciliary transport particle A
phototransduction
cellular response to light stimulus
photoreceptor activity
protein-chromophore linkage
G-protein coupled photoreceptor activity
biosynthesis of antibiotics
carbon metabolism
glycolytic process
glycolysis/gluconeogenesis
biosynthesis of amino acids
fatty acid degradation
fatty acid metabolism
PPAR signaling pathway
fatty acid metabolic process
valine, leucine, and isoleucine degradation
fatty acid beta-oxidation
Cep290, Arl6, Tub, Bbs5, Bbs12, Bbip1, Ift81, Arl13b, Prom1, Rp1, Kif19A, Tulp1, Sirt2, Cetn4
Ttc21a, Dync2h1, Bbs1, Bbs9, Bbs4, Ift122, Wdr35
Gnat1, Rgr, Cngb1, Rpgr, Gngt1, Guca1a, Opn1sw, Rcvrn, Rho, Rp1
Hk1, Psph, Acat1, Mdh1, Pkm, Pgk1, Ldha, Aldoa
Acadm, Acadvl, Cpt2, Acsl1, Acsl6
0 5 10 15

**Figure 3—Supplement 2. Global transcriptional changes in P4 NMNAT1 knockout retinas.** (**A**) schematic depicting RNA-sequencing experimental approach for P4 knockout and control retinas. (**B**) donut chart representing relative proportions of differentially upregulated and downregulated genes in P4 knockout retinas. (**C**) results of pathway overrepresentation analysis of differentially-expressed genes in P4 knockout retinas. n=5 biological replicates.

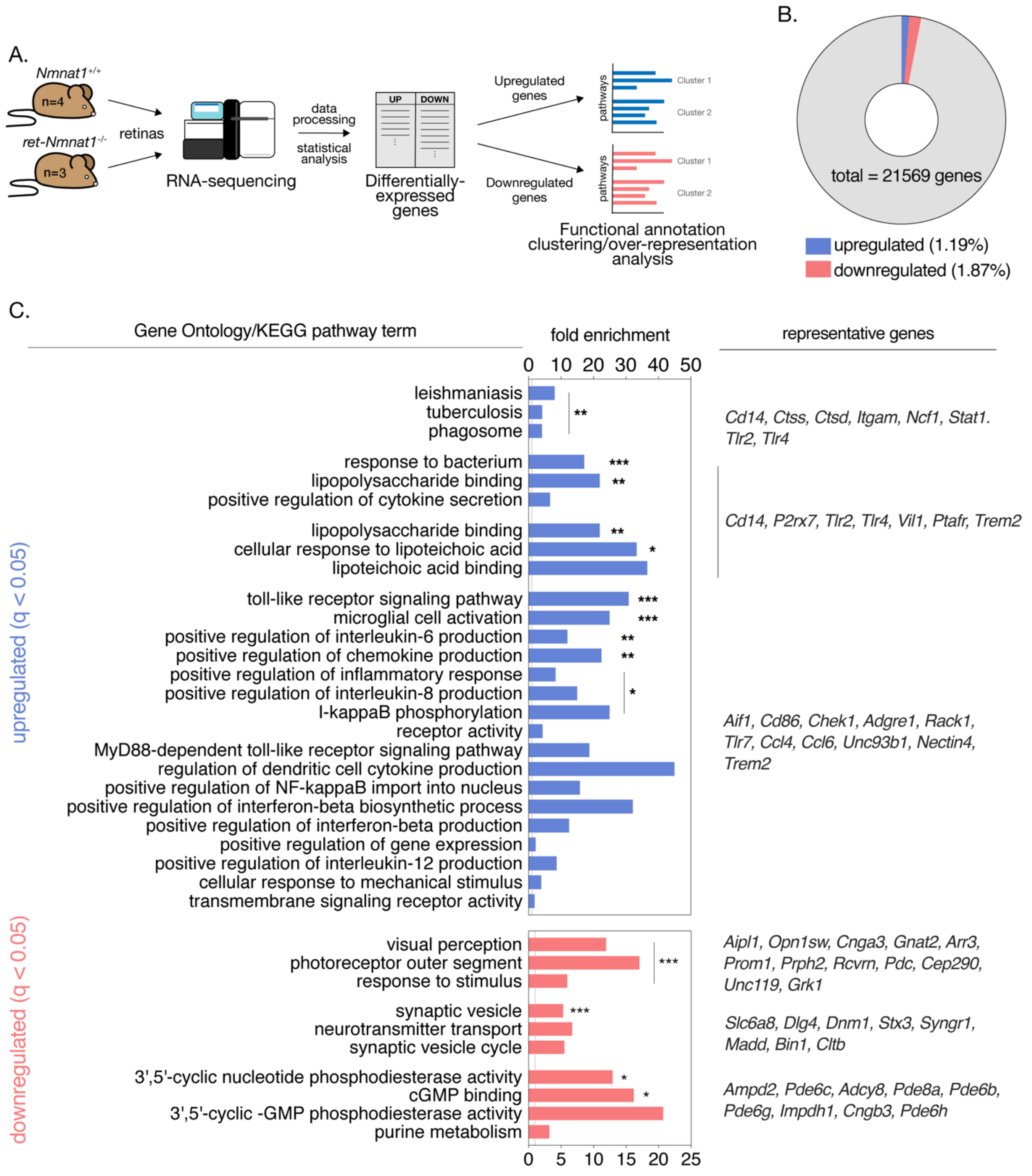


**Figure 3—Supplement 3. Global transcriptional changes in E18.5 NMNAT1 knockout retinas.** (**A**) schematic depicting RNA-sequencing experimental approach for E18.5 knockout and control retinas. (**B**) donut chart representing relative proportions of differentially upregulated and downregulated genes in E18.5 knockout retinas. (**C**) results of pathway overrepresentation analysis of differentially-expressed genes in E18.5 knockout retinas. n=4 biological replicates (one outlier removed from knockout group).

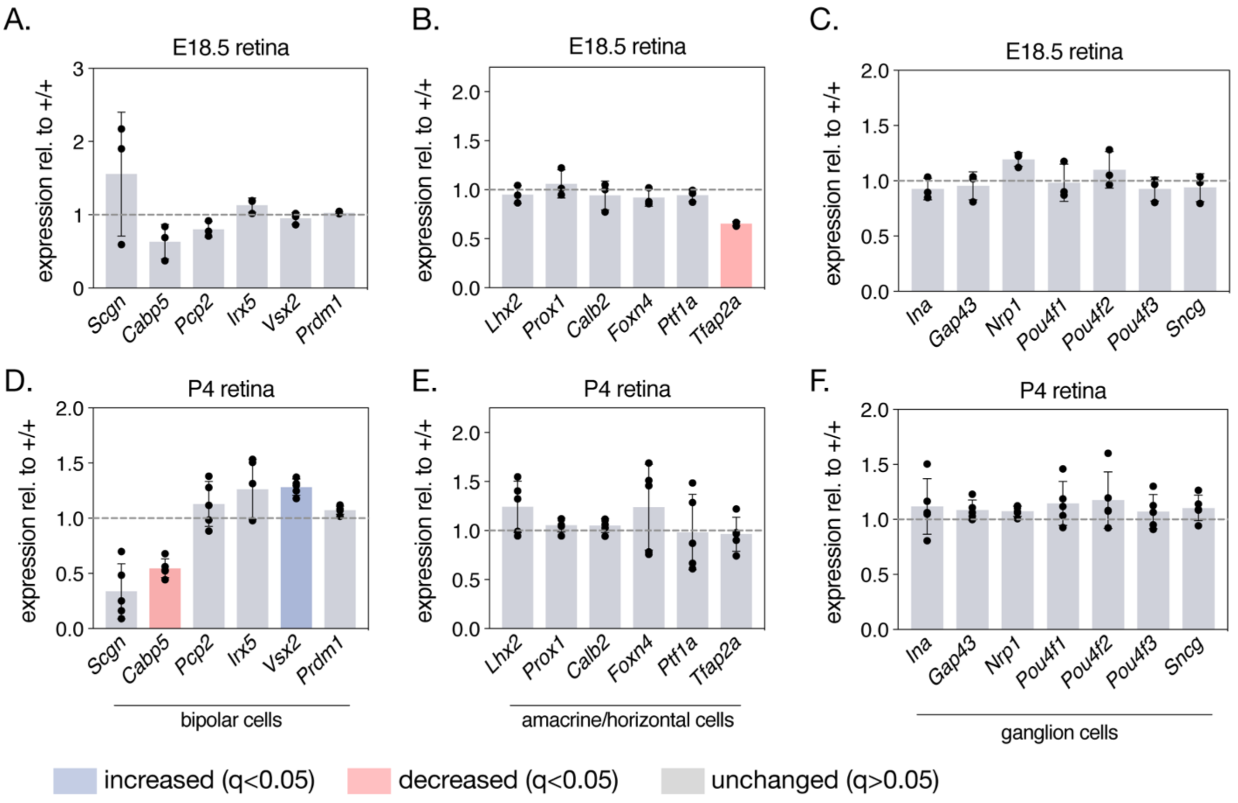


**Figure 3—Supplement 4. Expression of a collection of non-photoreceptor specific genes is largely unchanged in NMNAT1 knockout retinas.** Relative expression of indicated bipolar cell (**A**,**D**), amacrine and horizontal cell (**B**,**E**) and ganglion cell (**C**,**F**) genes in E18.5 (**A-C**) or P4 (**D-F**) knockout retinas as assessed by RNA-sequencing. Data are represented as mean ± SD. n= 3 biological replicates for (**A-C**), n=5 biological replicates for (**D-F**).

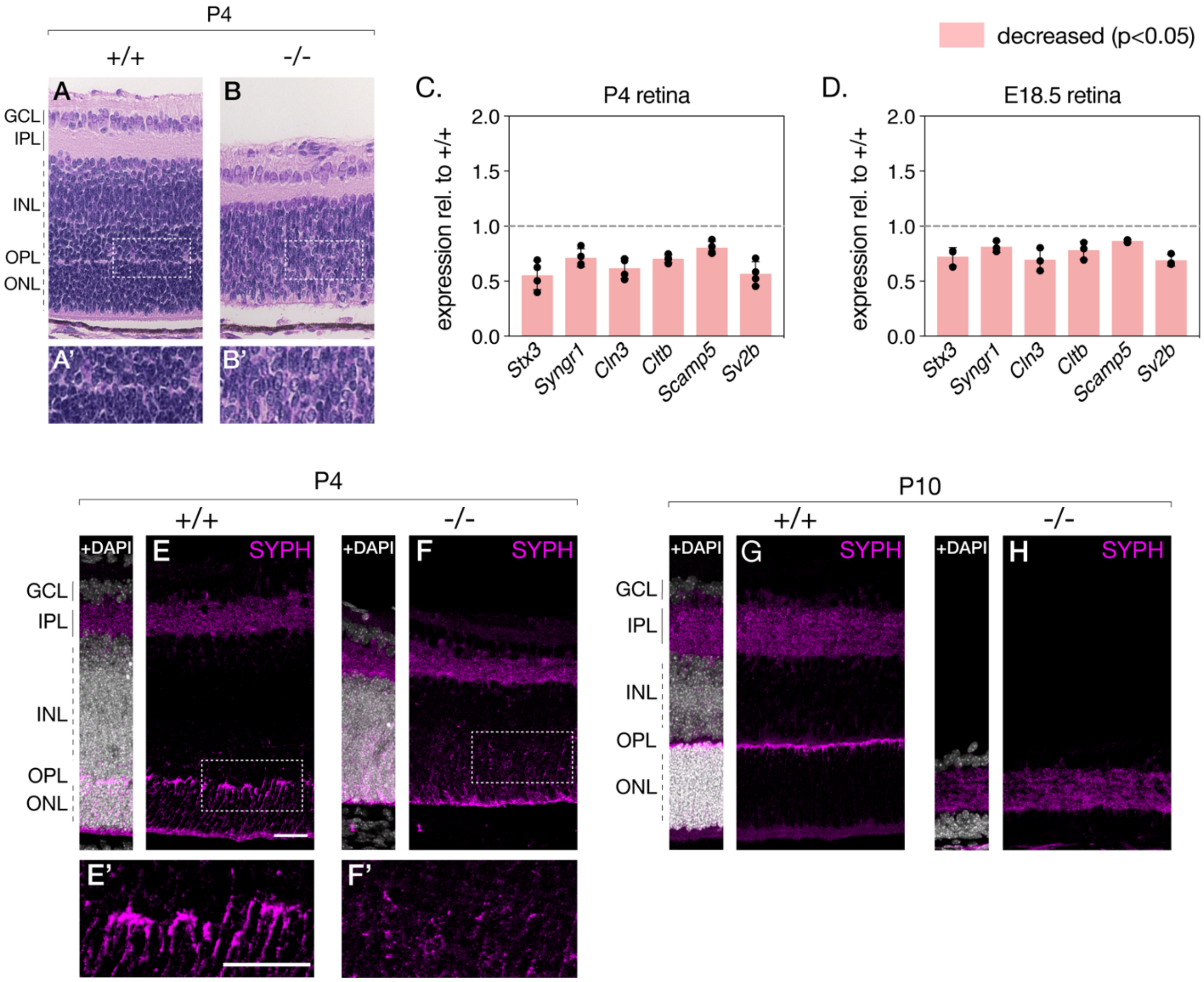


**Figure 3—Supplement 5. NMNAT1-loss during retinal development affects formation of the outer plexiform layer.** (**A**) representative hematoxylin and eosin (H&E)-stained retinal cross-sections from P4 knockout (**B, B'**) and control (**A, A'**) mice. Relative expression of indicated synaptic genes in P4 (**C**) and E18.5 (**D**) knockout retinas as assessed by RNA-sequencing are also shown. (**E-H**) representative retinal sections from knockout (-/-) and floxed littermate control (+/+) mice at the indicated ages labelled with antibodies against synaptophysin (SYPH). Data are represented as mean ± SD. n=5 biological replicates for (**C**), n=3 biological replicates for (**D**,**E-H**). Corresponding zoom panels are indicated with dotted rectangles. Scale bars, 30 μm.

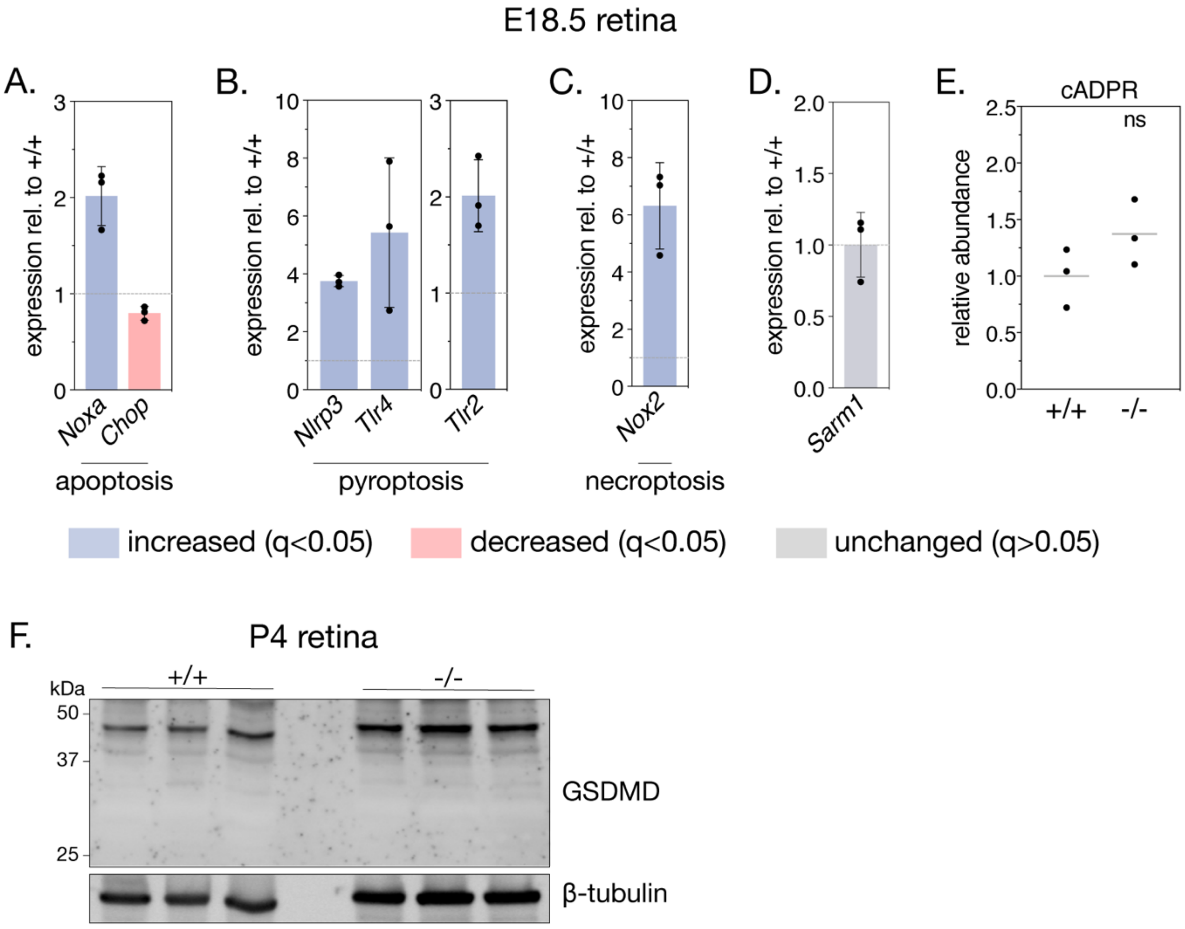


**Figure 4—Supplement 1. Deregulation of several cell death pathways in NMNAT1-null retinas preceding gross degeneration.** Relative expression of several apoptotic (**A**), pyroptotic (**B**), and necroptotic (**C**) genes in E18.5 knockout retinas as assessed by RNA-sequencing. (**D**) relative expression of *Sarm1* in E18.5 knockout and control retinas as assessed by RNA-sequencing. (**E**) relative abundance of cyclic-ADP-ribose (cADPR) in E18.5 knockout and control retinas as measured by mass spectrometry (grey bars represent means). (**F**) western blot of retinal lysate from P4 knockout and control retinas stained with an antibody against gasdermin D (GSDMD) and tubulin loading control. Data are represented as mean ± SD. n=5 biological replicates for (**A-D**), n=3 biological replicates for (**E**). Significance determined using DeSeq2 (see methods) (**A-D**) or Student's t-test (**E**).

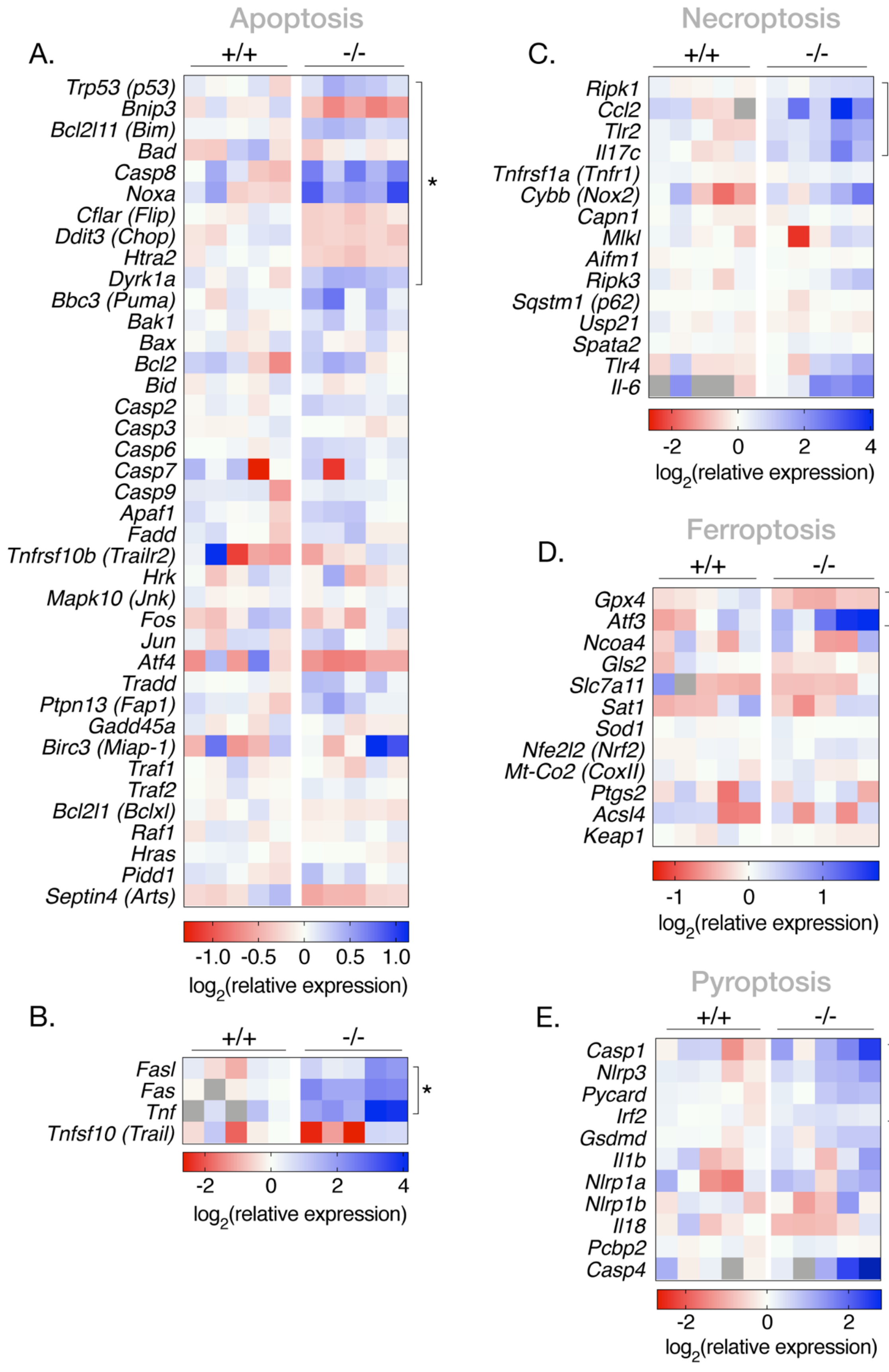
Apoptosis
A.
+/+
-/-
Trp53 (p53)
Bnip3
Bcl2l11 (Bim)
Bad
Casp8
Noxa
Cflar (Flip)
Ddit3 (Chop)
Htra2
Dyrk1a
Bbc3 (Puma)
Bak1
Bax
Bcl2
Bid
Casp2
Casp3
Casp6
Casp7
Casp9
Apaf1
Fadd
Tnfrsf10b (Trailr2)
Hrk
Mapk10 (Jnk)
Fos
Jun
Atf4
Tradd
Ptpn13 (Fap1)
Gadd45a
Birc3 (Miap-1)
Traf1
Traf2
Bcl2l1 (Bclxl)
Raf1
Hras
Pidd1
Septin4 (Arts)
*
-1.0 -0.5 0 0.5 1.0
log2(relative expression)
B.
+/+
-/-
Fasl
Fas
Tnf
Tnfsf10 (Trail)
*
-2 0 2 4
log2(relative expression)
Necroptosis
C.
+/+
-/-
Ripk1
Ccl2
Tlr2
Il17c
Tnfrsf1a (Tnfr1)
Cybb (Nox2)
Capn1
Mlkl
Aifm1
Ripk3
Sqstm1 (p62)
Usp21
Spata2
Tlr4
Il-6
*
-2 0 2 4
log2(relative expression)
Ferroptosis
D.
+/+
-/-
Gpx4
Atf3
Ncoa4
Gls2
Slc7a11
Sat1
Sod1
Nfe2l2 (Nrf2)
Mt-Co2 (CoxII)
Ptgs2
Acsl4
Keap1
*
-1 0 1
log2(relative expression)
Pyroptosis
E.
+/+
-/-
Casp1
Nlrp3
Pycard
Irf2
Gsdmd
Il1b
Nlrp1a
Nlrp1b
Il18
Pcbp2
Casp4
*
-2 0 2
log2(relative expression)

**Figure 4—Supplement 2. Transcriptional survey of cell death in the NMNAT1-null retina.** Heatmaps depicting log-scaled expression (relative to average of control samples) of genes related to apoptosis (**A,B**), necroptosis (**C**), ferroptosis (**D**), and pyroptosis (**E**) in P4 knockout and control retina as assessed by RNA-sequencing. Asterisks depict statistical significance as determined using DESeq2 (see methods). n=5 biological replicates.

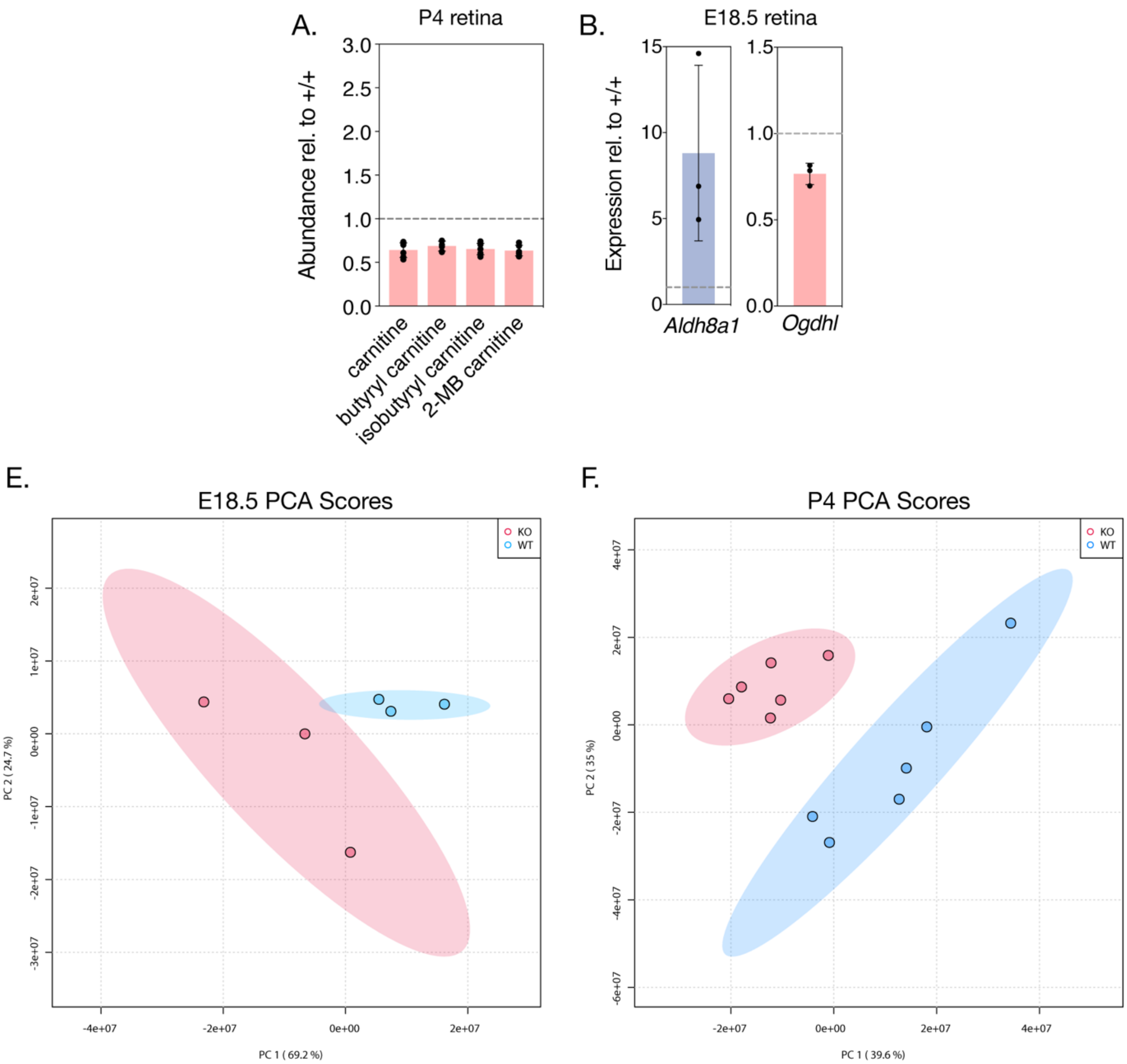


**Figure 5—Supplement 1. Additional metabolic changes in NMNAT1-null retinas.** (**A**) relative abundance of indicated carnitine species in P4 knockout retinas as assessed by mass spectrometry. (**B**) relative expression of *Aldh8a1* and *Ogdhl* in E18.5 knockout retinas as measured by RNA-sequencing. Principal component analysis (PCA) of E18.5 (**E**) and P4 (**F**) metabolomics data reveals expected clustering by genotype. Data are represented as mean ± SD. n=6 biological replicates for (**A**), n=5 biological replicates for (**B**).

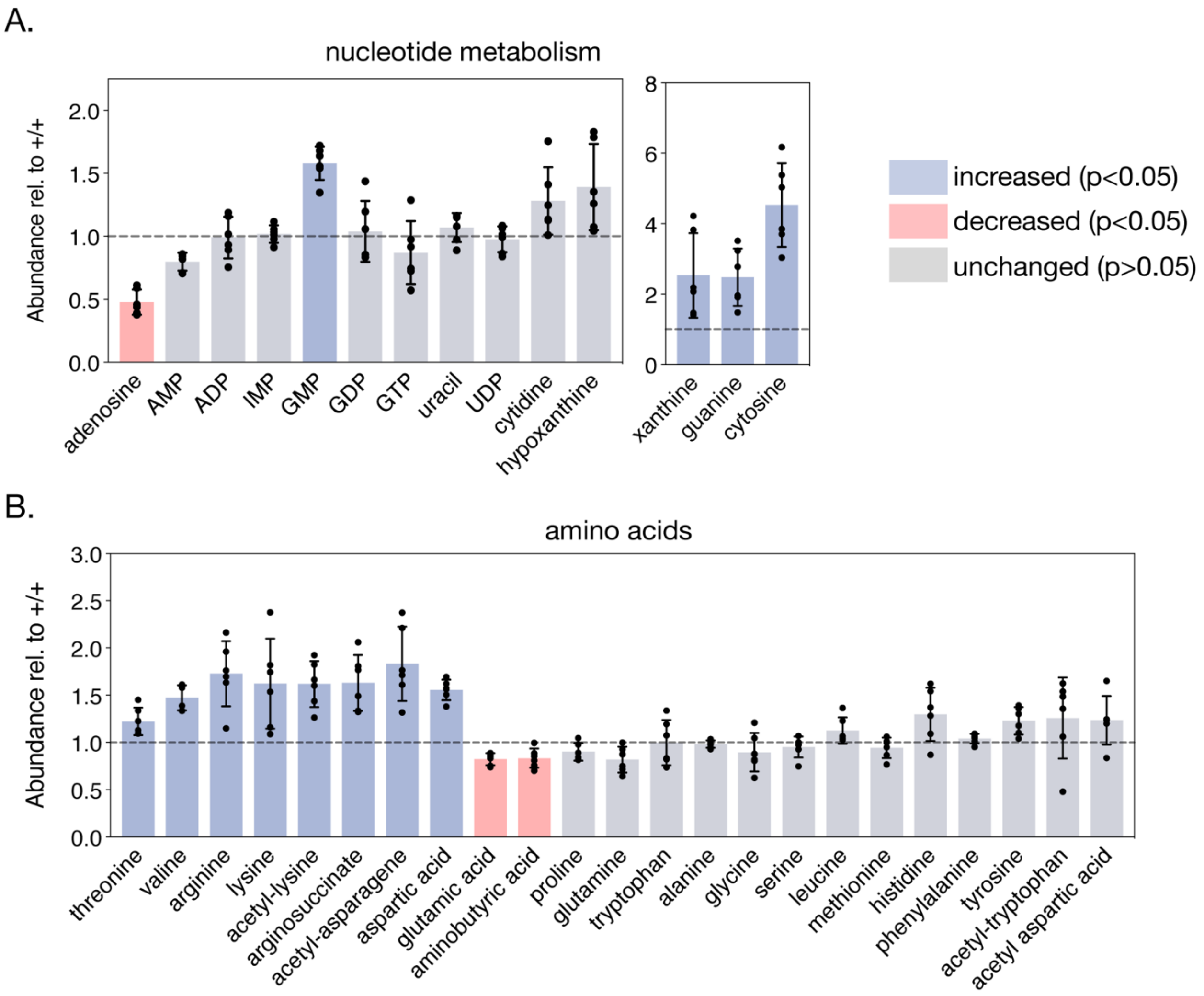


**Figure 5—Supplement 2. NMNAT1-associated changes in retinal nucleotide and amino acid metabolism.** (**A**) relative abundance of indicated nucleotide metabolites in P4 knockout retinas as assessed by mass spectrometry. (**B**) relative abundance of indicated amino acid metabolites in P4 knockout retinas as assessed by mass spectrometry. Data are represented as mean ± SD. n=6 biological replicates.

| Feature | Forward Primer Sequence | Reverse Primer Sequence |
|---|---|---|
| *Six3*-Cre | 5'-CCTGGAAAATGCTTCTGTCCG-3' | 5'-CAGGGTGTTATAAGCAATCCC-3' |
| *Nmnat1* 5' loxP | 5'-TCGGAGTGTATCCTTGGAGT-3' | 5'-ACCAAGCTTTCAGCACATGG-3' |
| *Nmnat1* 3' loxP | 5'-CCCAGTCACTAAGACATTCAA-3' | 5'-GACCCTCCTAGGCAAATATA-3' |
| *Nmnat1* (RT-qPCR) | 5'-CTTTTAACCCCATCACCAACATGC-3' | 5'-CCTTTCTTCTTGTACGCATCACC-3' |
| *Nmnat2* | 5'-CTTTTGTAGATGAGAACGCCAACC-3' | 5'-CCAACAATCACTTCCATATCTGCC-3' |
| *Nmnat3* | 5'-AAGACACCATCAGCCTCTGC-3' | 5'-CCAAGCCGAACTTCTCCACT-3' |
| *Cep290* | 5'-AAGGTACTGAGAAAATTGTTGCCG-3' | 5'-TGTAGTCTCTTCCCAGTTTCTTCG-3' |
| *Prom1* | 5'-TGAGACCCAAGATACCTTCAATGC-3' | 5'-AGACTATGATTTCTGGCTCCTTGG-3' |
| *Gnat1* | 5'-GGAGAAGAAGCTGAAAGAGGATGC-3' | 5'-AGAGTGTTGCCGTAGATGATGG-3' |
| *Rp1* | 5'-CAAGTTACCAGGAATCTCTCATCG-3' | 5'-TCTAAGGCCAAGTAATTCTCAGGG-3' |
| *Hmbs* | 5'-GTTTACCAAGGAGCTAGAAAACGC-3' | 5'-GTGAAAGACAACAGCATCACAAGG-3' |
| *Ppia* | 5'-GGATTTGGCTATAAGGGTTCCTCC-3' | 5'-GTTCTCATCCTCAAATTTCTCTCCG-3' |
| *Ywhaz* | 5'- GTTGTAGGAGCCCGTAGGTCATCG-3' | 5'- GCTTTCTGGTTGCGAAGCATTGGG-3' |

**Supplementary Table 1. Primer sequences for genotyping and RT-qPCR experiments.**

| Designation | Source or reference | Identifiers | Additional Information |
|---|---|---|---|
| Anti-NMNAT1 (rabbit polyclonal) | this study | N/A | WB: 1:1000 |
| Anti-β tubulin (mouse monoclonal) | Sigma | Cat#: T5201 | WB: 1:3000 |
| Anti-FLAG (mouse monoclonal) | Sigma | Cat#: F1804 | WB: 1:2000 |
| Anti-brn3a (mouse monoclonal) | Santa Cruz | Cat#: sc-8429 | IF: 1:150 |
| Anti-calretinin (rabbit polyclonal) | Millipore Sigma | Cat#: AB5054 (discontinued) | IF: 1:500 |
| Anti-CHX10 (mouse monoclonal) | Santa Cruz | Cat#: sc-365519 | IF: 1:50 |
| Anti-recoverin (rabbit polyclonal) | Sigma | Cat#: AB5585 | IF: 1:500 |
| Anti-rhodopsin (4D2) (mouse monoclonal) | R. Molday, Univ. British Columbia | N/A | IF: 1:500 |
| Anti-synaptophysin (rabbit monoclonal) | Thermo Fisher | Cat#: MA5-14532 | IF: 1:1000 |
| Anti-active caspase-3 (rabbit | R&D Systems | Cat#: AF835 | IF: 1:500 |
| 4′,6-diamidino-2-phenylindole (DAPI) | Thermo Fisher | Cat#: D1306 | IF: 1:2000 |
| goat anti-rabbit Alexa Fluor-568 | Invitrogen | Cat#: A-11011 | IF: 1:1000 |
| goat anti-mouse Alexa Fluor-488 | Invitrogen | Cat#: A-11001 | IF: 1:1000 |
| goat anti-rabbit Alexa Fluor 680 | Invitrogen | Cat#: A-21076 | WB: 1:50000 |
| goat anti-mouse DyLight 800 | Invitrogen | Cat#: SA5-10176 | WB: 1:50000 |

**Supplementary Table 2. Antibodies used in this study.**

| Metabolite | HMDB | Positive/ Negative mode | Q1 Mass (Da) | Q3 Mass (Da) | Declustering Potential (Volts) | Collision Energy (Volts) | Retention time (RT) |
|---|---|---|---|---|---|---|---|
| 1-Methyladenosine | HMDB0003331 | Pos | 282.1 | 150.1 | 48 | 35 | 1.85 |
| 2-Methylbutyroylcarnitine | HMDB0000378 | Pos | 246.2 | 85.1 | 100 | 43 | 1.3 |
| D-2-Hydroxyglutaric acid | HMDB0000606 | Pos | 149.1 | 77 | 100 | 43 | 1.37 |
| 3-Aminoisobutanoic acid | HMDB0003911 | Pos | 104.1 | 58 | 55 | 38 | 1.37 |
| 3-Hydroxybutyric acid | HMDB0000357 | Pos | 105.1 | 45 | 20 | 47 | 1.3 |
| Hydroxykynurenine | HMDB0000732 | Neg | 223 | 75 | -150 | -55 | 0.3 |
| p-Aminobenzoic acid | HMDB0001392 | Pos | 138.1 | 120 | 63 | 24 | 0.46 |
| 4-Hydroxyphenylpyruvic acid | HMDB0000707 | Neg | 179.1 | 97.1 | -40 | -17 | 4.08 |
| 4-Hydroxyproline | HMDB0000725 | Pos | 132.1 | 86 | 60 | 17 | 2.63 |
| N-Acetyl-L-aspartic acid | HMDB0000812 | Pos | 176.1 | 74 | 50 | 20 | 3.4 |
| Acetyl-CoA | HMDB0001206 | Pos | 810.1 | 303.1 | 116 | 46 | 3.98 |
| N-Alpha-acetyllysine | HMDB0000446 | Pos | 189.1 | 84.1 | 50 | 32 | 5 |
| N-acetyltryptophan | HMDB0013713 | Pos | 247.2 | 146 | 75 | 35 | 0.42 |
| N-Acetylasparagine | HMDB0006028 | Pos | 175.1 | 70.1 | 55 | 24 | 3.1 |
| N-Acetylglutamic acid | HMDB0001138 | Pos | 190.1 | 84.1 | 51 | 20 | 3.02 |
| cis-Aconitic acid | HMDB0000072 | Neg | 173 | 85 | -37 | -18 | 3.66 |
| Adenine | HMDB0000034 | Neg | 134 | 107 | -92 | -25 | 0.73 |
| Adenosine | HMDB0000050 | Pos | 268.1 | 136 | 95 | 38 | 0.79 |
| Adipic acid | HMDB0000448 | Neg | 145 | 83 | -50 | -17 | 3.14 |
| ADP | HMDB0001341 | Neg | 426 | 79.1 | -65 | -84 | 4.6 |
| Oxoglutaric acid | HMDB0000208 | Neg | 145 | 101 | -56 | -11 | 3.02 |
| L-Alanine | HMDB0000161 | Pos | 90 | 44 | 50 | 20 | 2.35 |
| Aminoadipic acid | HMDB0000510 | neg | 160.1 | 116.1 | -50 | -20 | 3.75 |
| Gamma-Aminobutyric acid | HMDB0000112 | Pos | 104.1 | 87 | 57 | 13 | 2.57 |
| Adenosine monophosphate | HMDB0000045 | Neg | 346 | 134 | -82 | -46 | 3.76 |
| L-Arginine | HMDB0000517 | Pos | 175.1 | 70.1 | 45 | 33 | 4.28 |
| Argininosuccinic acid | HMDB0000052 | Pos | 291.1 | 69.9 | 130 | 65 | 4.6 |
| Ascorbic acid | HMDB0000044 | Neg | 175 | 87 | -63 | -30 | 2.3 |
| L-Asparagine | HMDB0000168 | Pos | 133.1 | 70.1 | 55 | 24 | 2.94 |
| L-Aspartic acid | HMDB0000191 | Pos | 134.1 | 74 | 50 | 20 | 3.49 |
| Adenosine triphosphate | HMDB0000538 | Pos | 507.9 | 136.1 | 14 | 60 | 5.69 |
| Azelaic acid | HMDB0000784 | Neg | 187 | 97.1 | -73 | -25 | 2.48 |
| Betaine | HMDB0000043 | Pos | 118.1 | 58 | 166 | 56 | 1.67 |
| Biotin | HMDB0000030 | Neg | 243.1 | 200 | -84 | -23 | 1.62 |
| Butyrylcarnitine | HMDB0002013 | Pos | 232.2 | 85.1 | 72 | 30 | 1.5 |
| Cadaverine | HMDB0002322 | Pos | 103 | 77 | 190 | 25 | 1.02 |
| Cyclic AMP | HMDB0000058 | Neg | 328 | 134 | -87 | -32 | 2.28 |
| Carbamoyl phosphate | HMDB0001096 | Neg | 151 | 108 | -105 | -25 | 1.04 |
| L-Carnitine | HMDB0000062 | Pos | 163.1 | 85 | 100 | 15 | 2.6 |
| Carnosine | HMDB0000033 | Pos | 227.1 | 110.1 | 157 | 32 | 3.61 |
| Cyclic GMP | HMDB0001314 | Neg | 344 | 150 | -37 | -30 | 2.75 |
| Choline | HMDB0000097 | Pos | 104.1 | 60.1 | 95 | 37 | 1.51 |
| Citraconic acid | HMDB0000634 | Neg | 129 | 85 | -21 | -12 | 3.63 |
| Citric acid | HMDB0000094 | Neg | 191.1 | 87 | -73 | -25 | 2.39 |
| Citrulline | HMDB0000904 | Neg | 174.1 | 131.1 | -32 | -20 | 3.28 |
| Coenzyme A | HMDB0001423 | Pos | 768.1 | 261 | 34 | 47 | 4.3 |
| Creatine | HMDB0000064 | Pos | 132.1 | 90 | 170 | 17 | 2.66 |
| Creatinine | HMDB0000562 | Pos | 114 | 44 | 71 | 33 | 0.8 |
| L-Cystine | HMDB0000192 | Neg | 239 | 74 | -50 | -25 | 4.26 |
| Cytidine | HMDB0000089 | Pos | 244 | 112.1 | 50 | 30 | 1.39 |

| | | | | | | | |
|---|---|---|---|---|---|---|---|
| Cytosine | HMDB0000630 | Pos | 112 | 95 | 97 | 23 | 0.87 |
| Decanoylcarnitine | HMDB0000651 | Pos | 316.3 | 85.1 | 70 | 25 | 0.6 |
| Dihydroxyacetone phosphate | HMDB0001473 | Neg | 169 | 79 | -38 | 37 | 4.11 |
| Sphinganine | HMDB0000269 | Pos | 302.3 | 81 | 160 | 47 | 1.1 |
| Erythritol | HMDB0002994 | Pos | 123 | 80 | 40 | 33 | 0.38 |
| D-Erythrose 4-phosphate | HMDB0001321 | Neg | 199 | 97 | -160 | -15 | 4.1 |
| FAD | HMDB0001248 | Neg | 784.1 | 437.1 | 52.31 | 43 | 3.8 |
| Fructose 1,6-bisphosphate | HMDB0001058 | Neg | 339.1 | 79 | -56 | -75 | 8.74 |
| Glucose 1-phosphate | HMDB0001586 | Pos | 259 | 79 | -85 | -60 | 4.17 |
| Glucose 6-phosphate | HMDB0001401 | Neg | 259 | 79 | -46 | -62 | 4.11 |
| Guanosine diphosphate | HMDB0001201 | Pos | 444 | 152 | 65 | 26 | 5 |
| Geranyl-PP | HMDB0001285 | Neg | 313.1 | 185.1 | -58 | -45 | 0.27 |
| D-Glucose | HMDB0000122 | Neg | 179 | 89 | -50 | -15 | 2.07 |
| L-Glutamic acid | HMDB0000148 | Pos | 148.1 | 84.1 | 51 | 20 | 3.3 |
| L-Glutamine | HMDB0000641 | Pos | 147.1 | 84.1 | 45 | 23 | 3.03 |
| Glutaric acid | HMDB0000661 | Neg | 131 | 87 | -45 | -16 | 3.41 |
| Glycine | HMDB0000123 | Pos | 76 | 30.1 | 30 | 16 | 2.57 |
| Guanosine monophosphate | HMDB0001397 | Neg | 362.1 | 79 | -41 | -61 | 4.22 |
| Glutathione | HMDB0000125 | Neg | 306 | 143.1 | -61 | -26 | 3.8 |
| Oxidized glutathione | HMDB0003337 | Pos | 613 | 231 | 16 | 42 | 5.56 |
| Guanosine triphosphate | HMDB0001273 | Neg | 521.9 | 159 | -75 | -57 | 5.2 |
| Guanine | HMDB0000132 | Pos | 152 | 110 | 20 | 28 | 1.21 |
| Guanosine | HMDB0000133 | Neg | 282.1 | 150 | -67 | -33 | 1.64 |
| Heptadecanoic acid | HMDB0002259 | Neg | 269.1 | 135.1 | -77 | -35 | 1.05 |
| Hexanoylcarnitine | HMDB0000705 | Pos | 260.2 | 85.1 | 80 | 25 | 0.9 |
| Histamine | HMDB0000870 | Pos | 112 | 95 | 65 | 18 | 2.41 |
| L-Histidine | HMDB0000177 | Pos | 156.1 | 110 | 50 | 22 | 3.35 |
| L-Homoserine | HMDB0000719 | Pos | 120.1 | 74 | 37 | 16 | 3.04 |
| Hypotaurine | HMDB0000965 | Neg | 108.1 | 64 | -40 | -17 | 2.44 |
| Hypoxanthine | HMDB0000157 | Neg | 135 | 65 | -108 | -37 | 0.8 |
| Inosinic acid | HMDB0000175 | Neg | 347 | 79 | -118 | -86 | 4 |
| Inosine | HMDB0000195 | Neg | 267 | 135 | -123 | -30 | 1.1 |
| Isopentenyl pyrophosphate | HMDB0001347 | Neg | 245.1 | 79 | -60 | -50 | 2.7 |
| Isobutyryl-L-carnitine | HMDB0000736 | Pos | 232.1 | 85.1 | 70 | 30 | 1.5 |
| L-Kynurenine | HMDB0000684 | Neg | 207.1 | 144 | -58 | -33 | 1.59 |
| L-Lactic acid | HMDB0000190 | Neg | 89 | 43 | -60 | -17 | 1.13 |
| L-Leucine | HMDB0000687 | Pos | 132.1 | 86 | 60 | 16 | 1.65 |
| L-Lysine | HMDB0000182 | Pos | 147.1 | 84 | 38 | 22 | 4.23 |
| L-Malic acid | HMDB0000156 | Neg | 133 | 115 | -50 | -14 | 3.55 |
| Maleic acid | HMDB0000176 | Neg | 115 | 71 | -34 | -13 | 0.54 |
| Malonyl-CoA | HMDB0001175 | Pos | 854.1 | 347.1 | 75 | 45 | 4.97 |
| L-Methionine | HMDB0000696 | Pos | 150.1 | 61 | 27 | 43 | 1.65 |
| Methylmalonic acid | HMDB0000202 | Pos | 119 | 65 | 70 | 37 | 1.23 |
| myo-Inositol | HMDB0000211 | Neg | 179 | 87 | -105 | -24 | 2.9 |
| Tetradecanoylcarnitine | HMDB0005066 | Pos | 372.4 | 85.1 | 70 | 25 | 0.4 |
| Acetylglycine | HMDB0000532 | Neg | 116 | 74 | -37 | -14 | 1.92 |
| 1-Methylnicotinamide | HMDB0000699 | Pos | 137 | 78 | 110 | 37 | 1.63 |
| NAD | HMDB0000902 | Pos | 664 | 136 | 27 | 48 | 4.33 |
| NADH | HMDB0001487 | Neg | 664 | 397 | -7 | -46 | 3.91 |
| NADP | HMDB0000217 | Pos | 744 | 136 | 9 | 78 | 5.36 |
| NADPH | HMDB0000221 | Neg | 744 | 408 | -35 | -52 | 5.1 |
| Niacinamide | HMDB0001406 | Pos | 123 | 80 | 136 | 20 | 0.41 |
| Nicotinic acid | HMDB0001488 | Neg | 122 | 78 | -50 | -15 | 1.31 |

| | | | | | | | |
|---|---|---|---|---|---|---|---|
| Nicotinamide ribotide | HMDB0000229 | Pos | 335 | 123.1 | 40 | 25 | 4.7 |
| Nicotinamide riboside | HMDB00855 | Pos | 256.1 | 123 | 68 | 18 | 2.25 |
| L-Acetylcarnitine | HMDB0000201 | Pos | 204.1 | 85 | 71 | 30 | 1.86 |
| L-Octanoylcarnitine | HMDB0000791 | Pos | 288.3 | 85.1 | 100 | 25 | 0.7 |
| Ophthalmic acid | HMDB0005765 | Pos | 290.1 | 58 | 139 | 56 | 3.66 |
| Ornithine | HMDB0000214 | Pos | 133 | 70 | 47 | 23 | 4.3 |
| Oxalic acid | HMDB0002329 | Neg | 89 | 61 | -45 | -10 | 5.06 |
| Oxalacetic acid | HMDB0000223 | Neg | 131 | 87 | -30 | -11 | 3.79 |
| L-Palmitoylcarnitine | HMDB0000222 | Pos | 400.4 | 85.1 | 30 | 40 | 0.4 |
| Palmityl-CoA | HMDB0001338 | Pos | 1006.3 | 499.5 | 45 | 53 | 3.26 |
| Pantothenic acid | HMDB0000210 | Neg | 218.1 | 71 | -70 | -41 | 1.13 |
| L-Phenylalanine | HMDB0000159 | Pos | 166.1 | 120 | 50 | 18 | 1.37 |
| Phosphocreatine | HMDB0001511 | Neg | 210 | 79 | -42 | -49 | 4.42 |
| Phosphoenolpyruvic acid | HMDB0000263 | Neg | 167.1 | 79 | -38 | -20 | 3.95 |
| L-Proline | HMDB0000162 | Pos | 116.1 | 70.1 | 70,120 | 44 | 2.1 |
| Propionylcarnitine | HMDB0000824 | Pos | 218.1 | 85.1 | 60 | 24 | 1.9 |
| Pyroglutamic acid | HMDB0000267 | Pos | 130.1 | 84 | 100 | 20 | 2.02 |
| Quinic acid | HMDB0003072 | Neg | 191.1 | 85 | -60 | -33 | 2.54 |
| Riboflavin | HMDB0000244 | Pos | 377.1 | 243.1 | 14 | 29 | 1.07 |
| D-Ribulose 5-phosphate | HMDB0000618 | Neg | 229 | 79 | -70 | -45 | 3.43 |
| S-Adenosylmethionine | HMDB0001185 | Pos | 192.1 | 61 | 27 | 43 | 0.48 |
| L-Serine | HMDB0000187 | Pos | 106 | 60 | 40 | 22 | 2.83 |
| Spermine | HMDB0001256 | Pos | 203.1 | 129 | 33 | 26 | 1.9 |
| Stearoylcarnitine | HMDB0000848 | Pos | 428.5 | 85.1 | 70 | 25 | 0.3 |
| Succinic acid | HMDB0000254 | Neg | 117 | 73 | -54 | -16 | 2.92 |
| Taurine | HMDB0000251 | Pos | 126 | 108 | 200 | 15 | 1.8 |
| Thiamine | HMDB0000235 | Pos | 265 | 122.1 | 67 | 40 | 2.1 |
| L-Threonine | HMDB0000167 | Pos | 120.1 | 102 | 50 | 10 | 2.5 |
| Trimethylamine N-oxide | HMDB0000925 | Pos | 76 | 58 | 117 | 43 | 2.3 |
| Trigonelline | HMDB0000875 | Pos | 138 | 92 | 60 | 27 | 1.81 |
| L-Tryptophan | HMDB0000929 | Pos | 205 | 146 | 75 | 35 | 1.79 |
| L-Tyrosine | HMDB0000158 | Pos | 182.1 | 136 | 40 | 17 | 1.92 |
| Uridine diphosphate glucose | HMDB0000286 | Pos | 611 | 499 | 40 | 29 | 4.59 |
| Uracil | HMDB0000300 | Neg | 111 | 42 | | -37 | 0.53 |
| Uric acid | HMDB0000289 | Neg | 167 | 124 | -86 | -19 | 2.6 |
| Uridine | HMDB0000296 | Pos | 245 | 113 | 23 | 51 | 1.4 |
| L-Valine | HMDB0000883 | Pos | 118.1 | 72 | 60 | 14 | 1.76 |
| Xanthine | HMDB0000292 | Neg | 151 | 108 | -70 | -23 | 1.1 |
| Xanthosine | HMDB0000299 | Neg | 283.1 | 151 | -92 | -28 | 1.66 |
| Xanthurenic acid | HMDB0000881 | Neg | 204 | 160 | -67 | -19 | 1.15 |

**Supplementary Table 3. Mass spectrometry standards and parameters.**
